# Use of IFNγ/IL10 ratio for stratification of hydrocortisone therapy in patients with septic shock

**DOI:** 10.1101/502864

**Authors:** Rainer König, Amol Kolte, Olaf Ahlers, Marcus Oswald, Daniela Röll, George Dimopoulos, Iraklis Tsangaris, Eleni Antoniadou, Holger Bogatsch, Markus Löffler, Charles L. Sprung, Mervyn Singer, Frank Brunkhorst, Michael Oppert, Herwig Gerlach, Ralf A. Claus, Sina M. Coldewey, Josef Briegel, Evangelos J. Giamarellos-Bourboulis, Didier Keh, Michael Bauer

## Abstract

**Background:** Large clinical trials testing hydrocortisone therapy in septic shock have produced conflicting results. Subgroups may however benefit depending on their individual immune response.

**Methods:** We performed an exploratory analysis of the CORTICUS trial database employing machine learning to a panel of 137 variables collected from 83 patients (60 survivors, 23 non-survivors) including demographic and clinical measures, organ failure scores, leukocyte counts and circulating cytokine levels. The identified biomarker was validated against data collected from patients enrolled into a cohort of the Hellenic Sepsis Study Group (HSSG) (n=162) and two data sets of two other clinical trials. *Ex vivo* studies were performed on this biomarker to assess a possible mechanistic role.

**Results:** A low serum IFNγ/IL10 ratio predicted increased survival in the hydrocortisone group whereas a high ratio predicted better survival in the placebo group. Using this ratio for a decision rule, we found significant improvement in survival in the groups of patients being in compliance with the prediction rule (discovery set: OR=3.03 [95% Cl: 1.05-8.75], P=0.031, validation set: OR=2.01 [95% CI: 1.04-3.88], P=0.026). Applying the rule to two further, smaller datasets showed the same tendency. Mechanistic studies revealed that IFNγ/IL10 was negatively associated with pathogen load in spiked human blood. An *in silico* analysis of published IFNγ and IL10 values in bacteremic and non-bacteremic SIRS patients supported this association between the ratio and pathogen burden.

**Conclusion:** If confirmed prospectively, the IFNγ/IL10 ratio could be used as a rapidly available theranostic for use of hydrocortisone therapy in septic shock.

## Introduction

Though prospective, randomized, controlled multicentre trials have consistently reported faster shock resolution (1, 2), the utility of ‘low-dose’ hydrocortisone (HC) in patients with septic shock remains controversial. Whereas two French studies reported outcome benefit from a combination of hydrocortisone plus oral fludrocortisone (3, 4), the pan-European CORTICUS trial and the 5-country ADRENAL trial found no survival effect from hydrocortisone alone (2, 5). Possible explanations for this disparity included differences in mortality risk in the populations with a two-fold higher risk of mortality in the French control group (of (1)) compared to CORTICUS (61% *versus* 31%, respectively), and an increase in superinfections, variations in other aspects of clinical management, and genetic variations. Of note, a subset analysis of the ADRENAL trial indicated survival benefit from hydrocortisone in Australasian patients, no effect in British and Danish patients, and a trend to harm in patients enrolled in Saudi Arabia (2).

It is increasingly recognized that patients presenting in septic shock are hyper-inflamed yet at the same time immunosuppressed (6–8). Corticosteroids are traditionally considered to induce immune suppression via the glucocorticoid receptor (GR) and its repressive effect on pro-inflammatory transcription factors such as AP-1 and NFκB (9). Thus, patients in an overall state of immunosuppression may be potentially compromised by administration of an immunosuppressive drug. This argument is however complicated by an increasing evidence base implicating corticosteroids and GRs in immune-reconstitutive processes (10, 11). In human monocytes, corticosteroid treatment induced expression of innate immune-related genes, such as TLRs, and anti-inflammatory genes (10, 11). In macrophages and derived cell lines glucocorticoids induced a central component of the inflammasome (NLRP3) and, upon stimulation with endotoxin, induced secretion of pro-inflammatory cytokines such as tumor necrosis factor alpha (TNFα) (11). Furthermore, glucocorticoid-dependent NLRP3 induction resulted in sensitization of innate immune cells to extracellular ATP and thus an enhanced ATP-mediated secretion of pro-inflammatory cytokines following endotoxin stimulation (12). This immune-activating role of corticosteroids has been described as a response to acute stress enhancing the peripheral immune response, whereas chronic corticosteroid exposure leads to immune suppression (13, 14). These diverging effects of GCs support the need for biomarkers to guide their application. We applied machine learning to physiological and laboratory data from patients enrolled into a CORTICUS sub-study to determine a theranostic marker for hydrocortisone treatment. We found the ratio of serum interferon-γ (IFNγ) to interleukin-10 (IL-10) to identify specific sub-cohorts with increased and decreased survival upon treatment. We thus explored this predictive utility of this biomarker in further datasets of septic shock patients and performed *in vitro* mechanistic studies to explore the possible significance of this ratio.

## Materials and Methods

### The CORTICUS cohort

In addition to the standard CORTICUS protocol, the Berlin study group sampled blood for subsequent measurement of cytokines and other circulating inflammatory mediators from 84 patients in 13 participating sites. The study was approved by the local Ethics Committee (no: 153/2001). Written informed consent was obtained from patients, proxies or their legal representatives. Eligible patients were enrolled if they met the following inclusion criteria: clinical evidence of infection, evidence of a systemic response to infection, the onset of shock within the previous 72 hours and hypoperfusion or organ dysfunction attributable to sepsis. Notable exclusion criteria included an underlying disease process with a poor prognosis, life expectancy <24 hours, long-term immunosuppression, and treatment with long-term corticosteroids within the past 6 months or short-term corticosteroids within the past 4 weeks. Detailed eligibility criteria are given in Supplementary Table S1 and the original study (5). Patients were randomized to receive either *placebo* or 200 mg hydrocortisone (HC)/day for 5 days, followed by a tapering dose until day 11. Demographic and baseline characteristics were extracted from the CORTICUS database. 79 out of 83 (95%) patients received norepinephrine at baseline while none received epinephrine. Blood samples were taken directly before an ACTH stimulation test and administration of the study medication. Table S2 shows the timing of blood sampling relative to the onset of shock. Blood samples were collected on day 0, on day 2, on the morning of day 5 (end of full dose HC application), on day 12 (day after HC cessation), day 17 and 27. The soluble mediators interleukin-(IL)-6, −8, −10, −12 p70, IFNγ, TNFα, soluble TNF-receptor-I (sTNF-RI), soluble FAS (all OptEIA (™) Set Human, BD Biosciences, New Jersey, USA), and E-selectin (R&D, Minnesota, USA) were measured in serum, plasma, or culture supernatant by enzyme-linked immunosorbent assay (ELISA) according to the manufacturers’ instructions. This included calculating calibration and standard curves. All measurements were performed in duplicate. The cytokine and all other laboratory values were taken from the original study.

One patient was removed due to lack of cytokine data. For ten patients either IFNγ or IL10 were below the detection limit. Excluding these patients did not change the overall results (Supplementary Text S8). Of the remaining 83 patients, serum lactate values (pre-treatment) and temporal values were available in 53 and 41 patients, respectively. For full details, see Supplementary Text S1 and S2.

### The Hellenic Sepsis Study Group (HSSG) cohort and two further cohorts for validation

Validation was performed on data obtained by the Hellenic Sepsis Study Group from septic shock patients with community-acquired pneumonia or intraabdominal infection. This study included a prospective collection of clinical data and biosamples from patients admitted to 45 study sites in Greece. Patients were enrolled after written informed consent provided by themselves or their legal representatives. Detailed eligibility criteria are given in Supplementary Table S1. All enrolled patients had been reclassified into infection and sepsis using the Sepsis-3 classification criteria (15, 16). In HC-treated patients 200 mg/day HC had been administered for 7 days followed by gradual tapering. Secreted cytokines were measured using the LEGENDplex Human Inflammation Panel (13-plex) (BioLegend, San Diego, USA) according to the manufacturer’s instructions with half of the reagents volume and sample incubation at 4°C overnight. After quality control and discarding data from patients dying or being discharged on the day of admission, a total of 162 eligible shock patients (HC treatment: n=63, No HC treatment: n=99) were selected. If only one of the cytokines (IFNγ or IL10) was below the detection limit, the respective value was set as the detection limit. Excluding these samples from the analysis did not alter the findings (Supplementary Text S8). For further details, see Supplementary Text S3 and Table S3 for patient characteristics. Furthermore, we investigated serum IFNγ/IL10 of patients in the *placebo* arm of the randomized *placebo*-controlled, trial of Sodium Selenite and Procalcitonin-guided antimicrobial therapy in Severe Sepsis (SISPCT) (17). After propensity score matching, n=24 patients were included in the analysis (details about the study and statistics, see Supplementary Text S6). In addition, we analyzed serum IFNγ/IL10 of patients from an earlier small crossover study (18) (details about this study and the crossover scheme is given in Supplementary Text S7). In this study, the early arm got a comparable HC application as the HC arm of CORTICUS, and hence was also used for validating our marker. The cytokine values of the crossover study were taken from the original publication.

### *Ex vivo* whole blood culture experiments

To assess biological plausibility of the data-driven biomarker IFNγ/IL10, we performed *ex vivo* whole blood culture experiments in which blood of healthy donors was spiked with a broad range of bacterial products to simulate pathogen load. 200 µL of diluted (HBSS (1:1, V/V) heparinized whole blood obtained from healthy volunteers (n=5, male, 20-25 years) was stimulated with serial dilutions either of LPS (endotoxin of *E. coli B4:O111)* (Sigma Aldrich) or lysates from two *E. coli* isolates obtained from septic patients). *E. coli* lysates were created by sonification, following heat inactivation, and a serial dilution of the obtained fragment stock was performed. Following incubation (37°C, 18 hours, gentle agitation at 2 rpm) plasma supernatant was prepared by centrifugation (2500g at room temperature for 10 min), secreted cytokine levels were measured using the LEGENDplex Human Inflammation Panel (13-plex) (BioLegend), as described above. Comparison was made to vehicle controls.

### The rationale of the data analysis

We used the database consisting of 137 patient features (potential predictors) and 83 patients from the subcohort of the CORTICUS study population to perform an exploratory data analysis. This table also included the ratios of all cytokine combinations. We aimed to find a theranostic marker distinguishing HC responders (i.e. survivors) from non-responders (non-survivors). The complete list of available features is shown in Supplementary Table S4 and the workflow depicted in Supplementary Figure S1.

We sought the best predictor of 28-day survival for the placebo arm using one-level decision trees (stumps) calculated by a leave-one-out cross-validation scheme. The best decision trees were chosen by intelligent enumeration. Table S5 lists the results from all 41 cross-validation runs. Of these, the predictor and threshold, which was in most trees (i.e. the predictor IFNγ/IL10) was applied to the HC arm. In 39 out of 41 runs, the cutoff value was 0.95 and applied to the HC arm leading to the 39.8 percentile of all 83 samples. Patients whose IFNγ/IL10 ratio ranked among the first 39.8% patients were denoted “low-ratio patients”, the others “high-ratio patients”. When we weighted all non-surviving patients higher or lower, we observed a linear relation between the optimal percentile and the corresponding mortality rates. Hence, we calculated these optimal cutoffs to gain a calibration curve with which we determined the cutoff for the validation sets (see also Results and Discussion, details for the method are in Supplementary Text S2, Figure S2). All analyses were carried out in R (www.r-project.org) using custom scripts. Odds ratio and statistical significance calculations are described in Supplementary Text S2.

### *Ex vivo* whole blood culture experiments

To assess biological plausibility of the data-driven biomarker IFNγ/IL10, we performed *ex vivo* whole blood culture experiments in which blood of healthy donors was spiked with a broad range of bacterial products to simulate pathogen load. 200 µL of diluted (HBSS (1:1, V/V) heparinized whole blood obtained from healthy volunteers (n=5, male, 20-25 years) was stimulated with serial dilutions either of LPS (bacterial endotoxin of *E. coli B4:O111*) (Sigma Aldrich) or lysates from two *E. coli* isolates obtained from septic patients). *E. coli* lysates were created by sonification, following heat inactivation, and a serial dilution of the obtained fragments stock was performed. Following exposition (37°C, 18 hours, gently agitation 2rpm) plasma supernatant was prepared by centrifugation (2.500g, RT, 10 min), secreted cytokine levels were measured using the LEGENDplex Human Inflammation Panel (13-plex) (BioLegend) as described above compared to vehicle control.

## Results

### Patient characteristics

Patient characteristics are summarized in Table 1. Median plasma concentrations of soluble mediators and leukocytes were not significantly different between the arms at baseline (Supplementary Table S6). Age differed but was not a confounder (see Supplementary Text S4).

**Table 1.** Patient characteristics of the studied CORTICUS sub-cohort

|  | HC<br>(n=42) | Placebo<br>(n=41) |
| --- | --- | --- |
| Sex (male/ female, n) | 29/ 13 | 30/ 11 |
| Age [years] | 59.4 (22-87)* | 69.4 (43-88) |
| Height [cm] | 171 (150-188) | 172 (150-195) |
| Weight [kg] | 79.5 (50-130) | 77.1 (53-127) |
| Admission category (n) |  |  |
| - Medical | 1 | 0 |
| - Emergency surgery | 39 | 37 |
| - Elective surgery | 2 | 4 |
| SOFA score at inclusion | 10.4 (4-18) | 9.8 (5-17) |
| SAPS II score |  |  |
| - First 24 hours at ICU | 47.9 (22-80) | 48.0 (16-88) |
| - Last 24 hours before inclusion | 43.7 (13-77) | 47.2 (16-88) |
| Time: sepsis -> inclusion [hours] | 29 (3-71) | 31 (1-67) |
| ACTH Responder/ Non-Responder (n) | 26/ 16 | 29/ 12 |
| Survival (n) |  |  |
| - Day 28 (survivors/ non-survivors) | 31/ 11 | 29/ 12 |
| - ICU (survivors/ non-survivors) | 27/ 15 | 28/ 13 |
| - Hospital (survivors/ non-survivors) | 23/ 17 | 26/ 15 |
| ICU- stay [days] all patients | 27.5 (3-160) | 23.9 (3-89) |
| ICU- stay [days] survivors | 23.3 (5-73) | 26.3 (3-89) |
| Hospital stay [days] all patients | 51.3 (11-207) | 45.5 (3-135) |
| Hospital stay [days] survivors | 50.5 (15-128) | 53.9 (17-135) |

\* : $p < 0.05$ ; Range : minimum and maximum

### IFNγ/IL10 stratifies CORTICUS patients

Analysis of baseline characteristics was performed on 137 variables including demographic and clinical variables, Sepsis-related Organ Failure Assessment (SOFA) scores, lymphocyte counts, plasma protein concentrations of cytokines and patient blood stimulation experiments (Supplementary Table S4). We performed a leave-one-out cross-validation with one-level decision trees (using only one predictor at a time) to the *placebo* arm to discriminate between 28-day-survivors and non-survivors. This led to a high true positive rate (83%, Table 2a). In 95% of the cross-validation runs, the serum IFNγ/IL10 ratio (referred to as IFNγ/IL10 in the following) with the same threshold (39.8 percentile of IFNγ/IL10 from all patients) was selected by the algorithm as the best predictor. Patients which IFNγ/IL10 ratios ranked among the first 39.8% patients were denoted “low-ratio patients”, the others “high-ratio patients”. Upon applying this predictor to HC-treated patients, the reverse behaviour was observed with a high true negative rate i.e. a low IFNγ/IL10 indicated a high likelihood of survival (85%) (Table 2b). A significant interaction effect (p=0.0083) was observed between the IFNγ/IL10 ratio and the treatment (Supplementary Text S11, Figure S3). These observations were then compiled into a decision rule: no HC treatment if the ratio was high, and HC treatment if low. This decision rule yielded an odds ratio (OR) of survival of 3.03 [95% Cl: 1.05-8.75], p=0.031. Of note, neither IFNγ nor IL10 alone could allow this prediction. Our new treatment rule was applied to a validation dataset from the Hellenic Sepsis Study Group and to two smaller datasets from two other clinical trials.

**Table 2.**
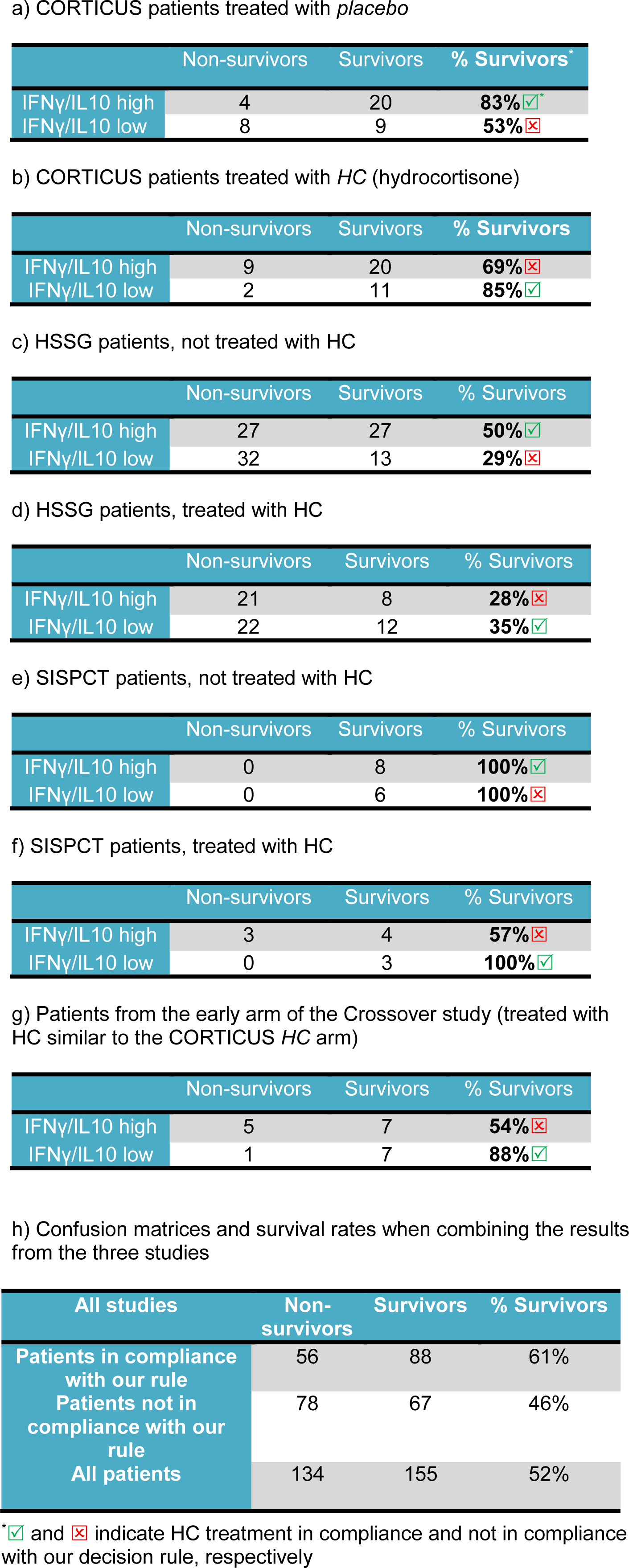
Survival rates according to high and low IFNγ/IL10

|  | Non-survivors | Survivors | % Survivors* |
| --- | --- | --- | --- |
| IFNγ/IL10 high | 4 | 20 | 83% |
| IFNγ/IL10 low | 8 | 9 | 53% |

b) CORTICUS patients treated with *HC* (hydrocortisone)
|  | Non-survivors | Survivors | % Survivors |
| --- | --- | --- | --- |
| IFNγ/IL10 high | 9 | 20 | 69% |
| IFNγ/IL10 low | 2 | 11 | 85% |

c) HSSG patients, not treated with HC
|  | Non-survivors | Survivors | % Survivors |
| --- | --- | --- | --- |
| IFNγ/IL10 high | 27 | 27 | 50% |
| IFNγ/IL10 low | 32 | 13 | 29% |

d) HSSG patients, treated with HC
|  | Non-survivors | Survivors | % Survivors |
| --- | --- | --- | --- |
| IFNγ/IL10 high | 21 | 8 | 28% |
| IFNγ/IL10 low | 22 | 12 | 35% |

e) SISPCT patients, not treated with HC
|  | Non-survivors | Survivors | % Survivors |
| --- | --- | --- | --- |
| IFNγ/IL10 high | 0 | 8 | 100% |
| IFNγ/IL10 low | 0 | 6 | 100% |

f) SISPCT patients, treated with HC
|  | Non-survivors | Survivors | % Survivors |
| --- | --- | --- | --- |
| IFNγ/IL10 high | 3 | 4 | 57% |
| IFNγ/IL10 low | 0 | 3 | 100% |

g) Patients from the early arm of the Crossover study (treated with HC similar to the CORTICUS *HC* arm)
|  | Non-survivors | Survivors | % Survivors |
| --- | --- | --- | --- |
| IFNγ/IL10 high | 5 | 7 | 54% |
| IFNγ/IL10 low | 1 | 7 | 88% |

h) Confusion matrices and survival rates when combining the results from the three studies
| All studies | Non-survivors | Survivors | % Survivors |
| --- | --- | --- | --- |
| Patients in compliance with our rule | 56 | 88 | 61% |
| Patients not in compliance with our rule | 78 | 67 | 46% |
| All patients | 134 | 155 | 52% |

\* and indicate HC treatment in compliance and not in compliance with our decision rule, respectively

### Validation based on patients from the Hellenic Sepsis Study Group (HSSG) and two further datasets

Table S3 summarizes demographics of the validation cohort. Applying IFNγ/IL10 to the group of HSSG patients, we observed a similar pattern as for the CORTICUS subgroup. High IFNγ/IL10 indicated distinct higher survival of the HC non-treated patients (50% *versus* 19%). In contrast, in the HC treated group we observed the opposite behaviour (28% *versus* 35%), yielding an odds ratio of OR=2.01 [95% CI: 1.04-3.88], P=0.026. As the survival rate of the HSSG patients was lower compared to CORTICUS, we had to adjust the threshold according to a calibration scheme based on the CORTICUS data explained in Methods. Figure 1 illustrates the potential of the theranostic marker to predict responders in both cohorts. Adjusting for an imbalance in co-morbidities between HC treated and non-treated patients by propensity score matching led to the same result (Supplementary Text S3).

**Figure 1.**
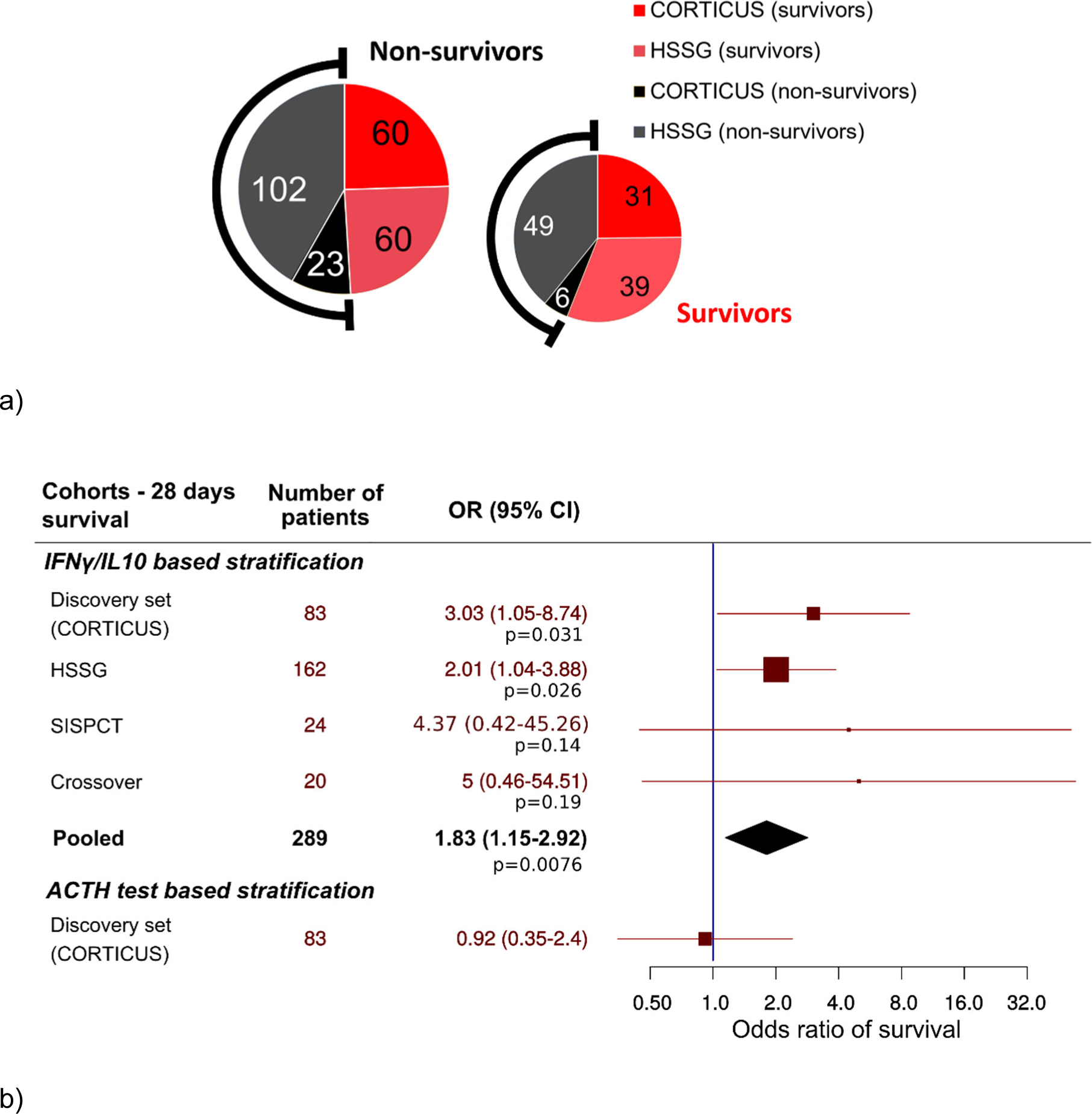
Pie chart and Forest plot for outcome in CORTICUS and HSSG cohorts depending on hydrocortisone administration according to the theranostic marker. (a) Survivors (red to light red) and non-survivors (black to grey) of all patients (left) and of patients being in compliance with the decision rule (right), patient numbers are given in the pies. Areas are proportional to number of patients. Treatment according to the decision rule was associated with an absolute risk reduction of 0.14 [95% CI: 0.03-0.26]. (b) Forest plot of the odds ratios for the sub-cohorts of CORTICUS (discovery set), HSSG (validation set) regarding IFNγ/IL10 prediction rule, SISPCT, the Crossover study, the combination of all cohorts, as well as the ACTH test (for CORTICUS subcohort) are given (details are provided in Supplementary Text S10). For SISPCT a pseudo-count was added to each field preventing division by zero.

Furthermore, we investigated serum IFNγ/IL10 of patients in 24 (propensity score matched) patients from the *placebo* arm of the randomized *placebo*-controlled, trial of Sodium Selenite and Procalcitonin-guided antimicrobial therapy in Severe Sepsis (SISPCT) (17) (details about the study and statistics, see Supplementary Text S6). All patients survived (n=3) if treated according to the rule, compared to 77% if not treated according to the rule. In addition, we analyzed serum IFNγ/IL10 of patients from an earlier small crossover study (18) in which the early arm got a comparable HC application as the HC arm of CORTICUS, and hence was also used for validating our marker (details about this study and the crossover scheme is given in Supplementary Text S7). In line to the results from CORTICUS, HSSG and SISPCT, low IFNγ/IL10 indicated good survival (88% survivors), whereas high IFN/IL10 was an indicator for considerably worse outcome (57% survivors), for HC treatment the survival rates between IFNγ/IL10 high and low were comparable (Table 2e). Due to the small sample sizes the results from SISPCT and the crossover study failed to achieve significance. In summary, the investigated patients from all studies evidenced IFNγ/IL10 as a potential theranostic marker for HC application in septic shock.

### Time courses of serum lactate and norepinephrine requirement reflect hemodynamic stabilization in patients treated in compliance with the decision rule

High serum lactate levels have been demonstrated to indicate severity of metabolic derangements and increased mortality in sepsis (15). Median initial lactate was 1.89 mmol/L in patients with high compared to 2.89 mmol/L in patients with low IFNγ/IL10 suggesting that IFNγ/IL10 associated with disease severity (boxplots, see Supplementary Figure S4). Of note, although the lactate levels correlate inversely with IFNγ/IL10 at baseline, serum lactate itself performed worse as a theranostic marker. Notably, we identified significant lactate decrease specifically in the group of patients, which were treated in compliance with the decision rule (Figure 2 a-d). Hemodynamic stabilization was also supported by the reduction of norepinephrine (NE) requirement (P=2.0e-05) specifically in the group of patients in compliance with the decision rule (Figure 2 e-h). In summary, time series of serum lactate and NE requirement reflect better recovery of septic shock in patients treated in compliance with the proposed theranostic marker.

**Figure 2.**
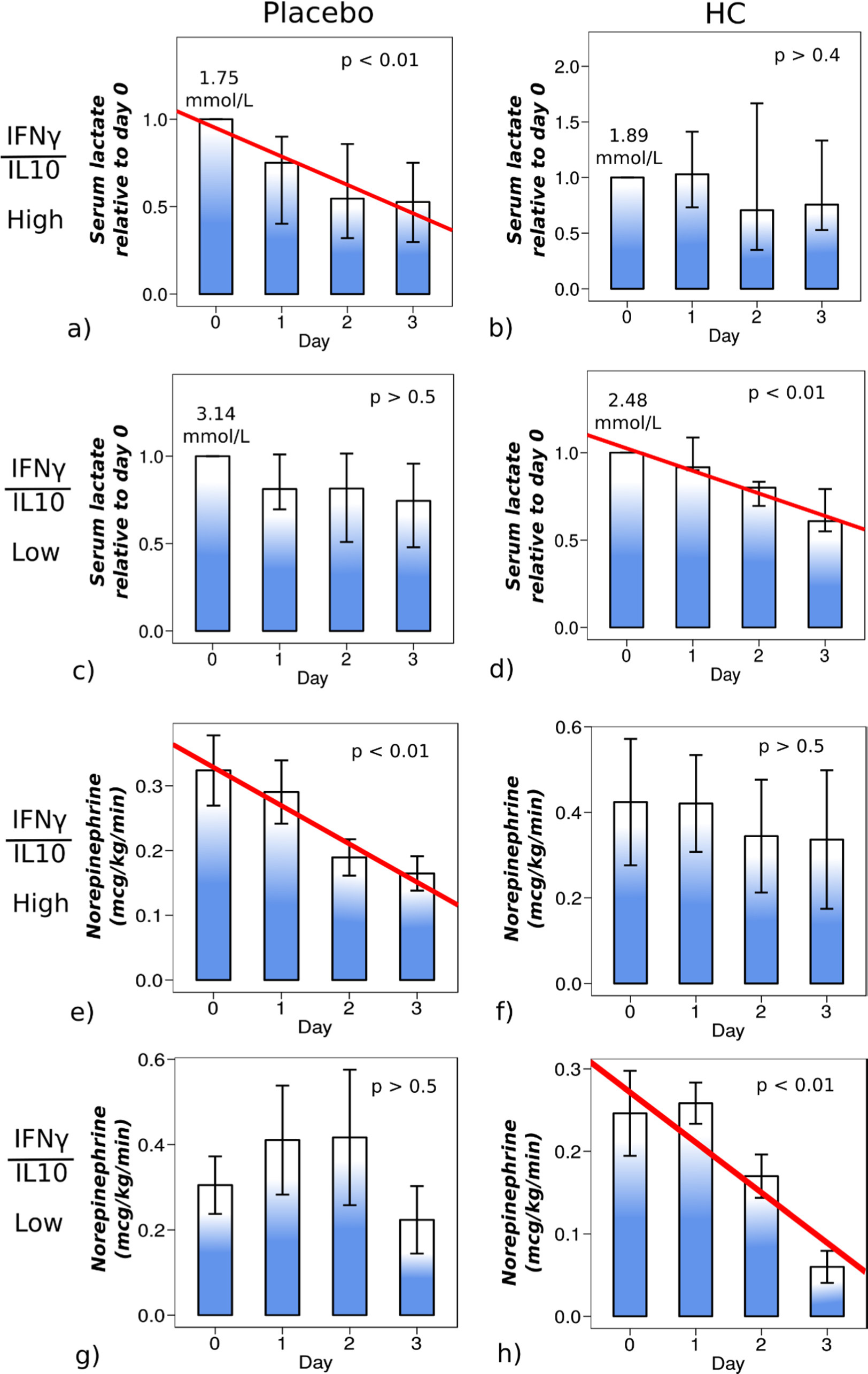
Time courses of serum lactate and norepinephrine consumption depending on hydrocortisone administration and IFN/IL10 ratio. (a) - (d) Normalized lactate levels, i.e. values for days 1-3 relative to day 0 for each patient are presented. Absolute serum lactate levels and p-values for the change over time in all available patients of the corresponding sub-group are reported in each panel. Patients in panels (a) and (d) reflect those in compliance with the treatment rule. For patients with high IFNγ/IL10, serum lactate decreased significantly in the *placebo* arm (P<0.01 n=11) (a), while its time course was rather heterogenous in the *HC* arm (n=13) (b). Among the low-ratio patients, there was a tendency of decrease in the *placebo* arm, however, not significant (n=12) (c), in contrast to a significant decrease in the *HC* arm (P<0.01, n=5) (d). (e) - (h) Norepinephrine consumption for days 1-3 is presented. Patients in panels (e) and (h) reflect those in compliance with the treatment rule. For patients with high IFNγ/IL10, NE consumption decreased significantly in the *placebo* arm (P<0.01 n=21) (e), while its time course was rather heterogenous in the HC arm (n=29) (f). Among the low-ratio patients, there was a heterogeneous trend in the placebo arm (n=20) (g), in contrast to a significant decrease in the HC arm (P<0.01, n=21) (h). Putting the compliant arms together (e and h), the rate of decrease over time was highly significant (p < 2e-05), but not significant in the non-complaint arm (panel f) and g), p > 0.5).

### IFNγ/IL10 as an indicator of pathogen load

In light of published evidence associating IFNγ and IL10 with the severity of parasitic and tuberculosis infection, we investigated whether IFNγ/IL10 reflects the pathogen burden of immune cells when challenged with typical pathogens found in sepsis. We spiked blood from healthy donors with *E. coli* fragments from clinical isolates or endotoxin across a wide range of concentrations, mimicking the immunological load. As expected, the higher the load, the higher the immune response and hence the concentration of IFNγ and IL10 in the supernatant (Figure 3a, b). Remarkably, we observed the inverse behaviour for the ratio, i.e. a high IFNγ/IL10 was observed for low pathogen concentrations and *vice versa* with “on-off” kinetics (Figure 3c). Results for a second *E. coli* isolate or endotoxin were similar (Supplementary Figure S5). To associate IFNγ/IL10 to the bacterial load in critical ill patients, we compared publically available data of patients with and without bacteremia. Matera *et al.* (19) investigated 52 patients (39 survivors and 13 non-survivors) with signs of SIRS at hospital admission. Only 4% of them were in septic shock. 28 of 52 were diagnosed with bacteremia. In line with our *ex vivo* observations, a distinctively higher ratio (1.6 ± 0.77) for IFN/IL10 in SIRS patients was observed compared to patients with bacteremia (0.8 ± 0.88, details: Supplementary Text S9). Healthy volunteers showed the highest IFNγ/IL10 (2.80 ± 1.22) of all groups investigated by Matera *et al*‥ In summary, IFNγ/IL10 inversely correlates with the immunological load of infection in an *in vitro* system and when comparing bacteremic and non-bacteremic critically ill patients.

**Figure 3.**
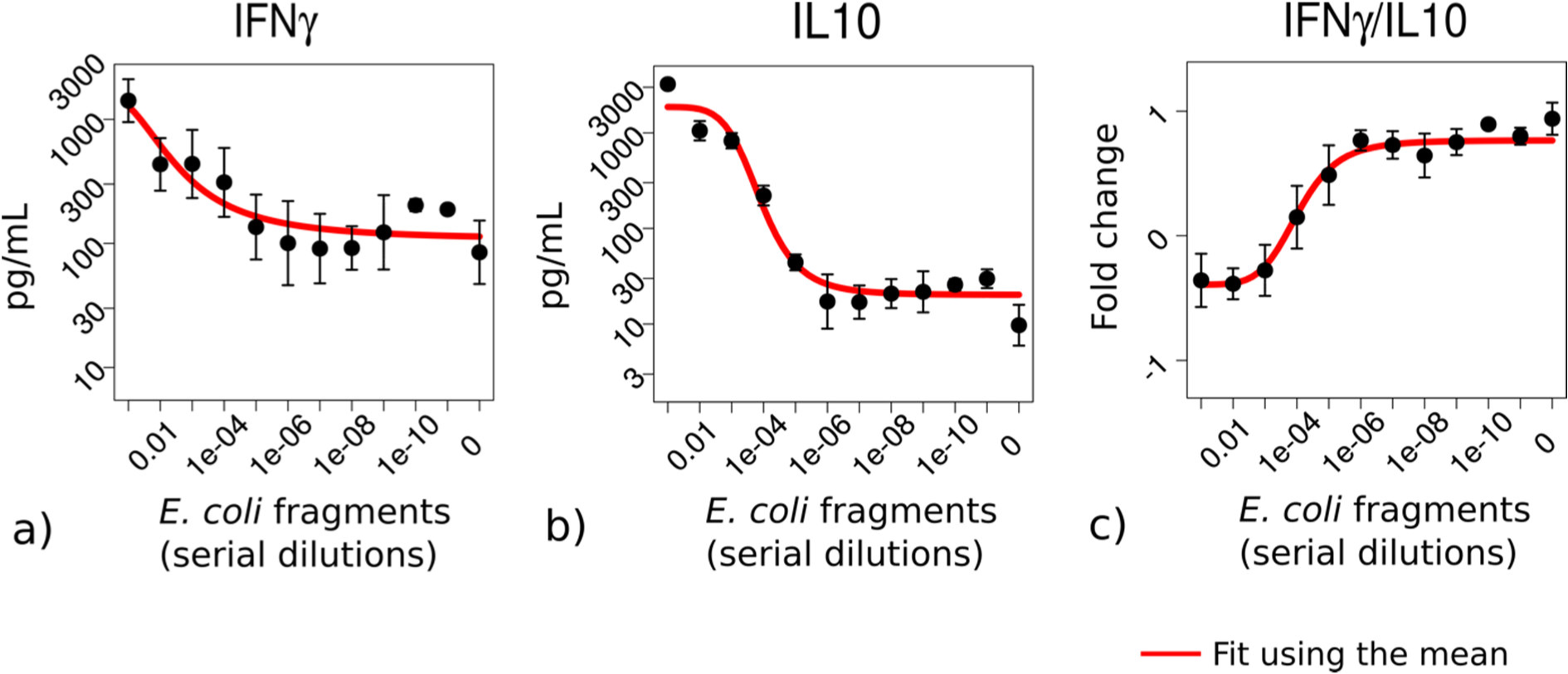
IFNγ/IL10 reflects the immunological load of immune cells *ex vivo*. Whole blood from five healthy donors was challenged with varying dilutions of *E. coli* fragments mimicking the bacterial burden on the immune system in sepsis. IFNγ (a) and IL10 (b) were elevated with increasing bacterial load while their ratio (c) showed the opposite behaviour, i.e. a high load was associated with a lower ratio.

### The corticotropin test and IFNγ/IL10 correlation to the severity of septic shock

Poor response to corticotropin stimulation is no longer recommended for HC treatment (3). We tried several machine learning concepts to stratify for treatment using serum HC baseline and after corticotropin stimulation. In line to results of the original study (5), the corticotropin test failed to predict response to treatment (Figure 1b, Supplementary Text S10, Table S7). We further investigated if IFNγ/IL10 correlates with the severity of septic shock. However, IFNγ/IL10 did neither correlate with mean arterial pressure nor with the SOFA score in the investigated CORTICUS cohort.

## Discussion

Starting with a larger panel of predictors from septic shock patients treated with HC or *placebo* in CORTICUS, we identified the ratio of serum IFNγ and IL10 as a promising biomarker for HC treatment decision. The decision rule was based on data derived from 83 patients. We validated our results by applying the marker and threshold to three other, unseen datasets to validate the potential of the theranostic marker. As in particular the survival rate of the HSSG cohort was very different to CORTICUS, this threshold needed to be adjusted. This was performed by a calibration curve calculated from the CORTICUS dataset mimicking higher death rates in CORTICUS. This led to a positive correlation between the optimal threshold and the mimicked death rate (details, see Supplementary Text S2), i.e. the higher the threshold, the more patients are in a worse condition at baseline (higher incidence of death), and particular these patients were observed to benefit from HC. Albeit the ratio was identified by machine learning and supported by subsequent *in vitro* experiments, there is published evidence for its plausibility as a biomarker for life-threatening infections. IFNγ and IL10 have been suggested as biomarkers for parasitic (20) and other infections, such as tuberculosis (21). McIlleron *et al*. reported the time course of IFNγ indicating response to treatment in pulmonary tuberculosis (22). However, for diagnosis of pulmonary tuberculosis, IFNγ alone has limitations with relation to sensitivity (23). In turn, high IL10 was observed with mycobacterial persistence (24) suggesting that in particular the ratio of these cytokines might reflect the immunological burden of infection. In line, it has been shown that the ratio of IFNγ and IL10 correlates with the disease severity of tuberculosis and may differentiate pulmonary from extrapulmonary forms (25). To elucidate the functional implications of low and high IFNγ/IL10, we performed blood culture assays. Indeed, we observed the ratio to be associated with the pathogen load in *ex vivo* blood culture experiments. This was supported by data from Matera *et al.* (19) when regarding critically ill patients with bacteremia (higher bacterial load) and without bacteremia (no or non-detectable bacterial load).

Synoptically, IFNγ/IL10 is associated with pathogen load and thus, severity of sepsis. Metabolically, high serum lactate indicates severity of cellular disturbances (15) and is associated with poor outcome (26). Consistent with this, patients with low IFNγ/IL10 showed high serum lactate. Furthermore, stratifying patients in compliance with the proposed decision rule, we observed a considerable decrease over time in serum lactate specifically in the group of patients, which were treated in compliance with the theranostic marker.

IFNγ has been consistently documented to activate cells of adaptive immunity (27) and IL10 as a suppressor of innate immune responses and inflammation (28, 29). According to this, high IFNγ/IL10 reflects increased activated adaptive *and* innate immunity. In line, we observed *ex vivo* that blood challenged with bacterial fragments or LPS, displayed low IFNγ/IL10 at high loads, and *vice versa*. Transferring these observations to septic shock patients with low IFNγ/IL10, we speculate that HC treatment yielded a better outcome in HC-responders because HC may allow the highly loaded immune system more pace for recovery. This is evidenced, by the reduction of serum lactate and norepinephrine requirements as measures of recovery particularly in the subgroup of very sick patients.

The concern about side effects of corticosteroids such as infections in patients with less severe septic shock stipulated more restrictive recommendations by the Surviving Sepsis Campaign. HC application is currently recommended for patients not responding to adequate fluid resuscitation and vasopressor therapy (30, 31). It seems intuitively promising to include an immune biomarker, such as IFNγ/IL10 reflecting the status of the patients’ immune system. Consistent with this concept, Bentzer *et al.* studied patients from the VASST trial investigating corticosteroid treated *versus* non-treated (only vasopressin or catecholamine vasopressors). They identified a signature of three cytokines (IL3, IL6, CCL4) suggesting response to corticosteroid treatment (32). However, these results are based on a study, which was not randomized, blinded or protocolized according to corticosteroid treatment. Further, Bentzer *et al.* did not distinguish between vasopressin and catecholamine treatment, and they didn’t elaborate explaining how these three cytokines interact with corticosteroid treatment.

Applying our decision rule successfully across a broad range of studies suggests the marker to be generic and rather independent from specific assays, probably reflecting the on-off kinetics of the quotient.

### Limitations and strengths

The relatively small sample sizes along with the high dimensional data sets obtained imply that our results must be interpreted with caution and requires prospective validation. A further limitation of our study was that the use of corticosteroids was not randomized in the HSSG and the SISPCT study. Strengths of our study were that we used well-phenotyped cohorts of patients with septic shock, that our marker showed very similar results across all studies, and that we not only showed the potential clinical relevance but also give a reasonable functional reasoning why this marker may support HC application by investigating *ex vivo* blood culture experiments.

## Conclusions

We identified the ratio of serum cytokines IFNγ and IL10 as a theranostic marker for hydrocortisone treatment in septic shock. An accompanying study into the mechanism suggests that this ratio enables the host to sense the pathogen load in an “on-off” fashion.

## Abbreviations

ACTH: Adrenocorticotropic hormone
ADRENAL: Adjunctive corticosteroid treatment in critically ill patients with septic shock
CI: Confidence interval
CORTICUS: Corticosteroid therapy of septic shock
HC: Hydrocortisone
HSSG: Hellenic sepsis study group
NE: Norepinephrine
OR: Odds ratio
SISPCT: Sodium selenite and procalcitonin guided antimicrobial therapy in severe sepsis
SOFA: Sequential organ failure assessment
VASST: Vasopressin and septic shock trial

## Declarations

### Acknowledgements

We thank Anne Goessinger for her outstanding support of the CORTICUS sub-study.

## Availability of data and materials

Most of the data is provided in the main text or supplementary material. The other datasets used or analyzed are available from the corresponding author on reasonable request.

## Authors’ contributions

RK, AK, MO, MB conceptualized and designed the study. MO, RK and AK developed the methodology. AK, MB, MS, JB, DK, DR and RK analyzed and interpreted the data. Administrative, technical, or material support was given by OA, DR, GD, IT, EA, HB, ML, CLS, MS, FB, MO, HG, RAC, SMC and JB. The study was supervised by RK, DK and MB. All authors read and approved the final manuscript.

## Ethics approval and consent to participate

The CORTICUS trial was a multicenter study, the protocol was approved by the ethics committee at each of the 52 participating intensive care units. In addition to the standard CORTICUS protocol, the Berlin study group sampled blood for subsequent measurement of cytokines and other circulating inflammatory mediators. This was approved by the local ethics committee (no: 153/2001). Written informed consent was obtained from patients, proxies or their legal representatives. The HSSG cohort represents a prospective collection of clinical data and biosamples in 45 study sites in Greece. The study protocol was approved from the ethics committees of all participating hospitals. Patients were enrolled after written informed consent provided by themselves or by first-degree relatives if patients were unable to consent. The Placebo-Controlled Trial of Sodium Selenite and Procalcitonin Guided Antimicrobial Therapy in Severe Sepsis (SISPCT) was a multicenter clinical trial. It was conducted in 33 multidisciplinary intensive care units across Germany. The study protocol was approved by the ethics board of Jena University Hospital. Written informed consent was obtained from all patients or their legal representatives. Among these, 109 patients were included in the Munich (Ludwig-Maximilians-University, LMU) sub-study for which cytokine measurements for this study was performed using their blood samples (ethics votum amendment EudraCT: 2007–004333-42). The study protocol for the Crossover study was approved by the institutional (Charite, Berlin) ethics committee.

## Consent for publication

Not applicable.

## Main Legends

## List of supplementary material

The supplementary material is compiled in one file denoted as Supplementary material.

