## Supplementary material for "Use of IFNγ/IL10 ratio for stratification of hydrocortisone therapy in patients with septic shock"

### Content

Supplementary Text ... 3
Text S1. Blood samples and serum assays of the CORTICUS Berlin sub-cohort ... 3
Text S2. Calibration of the IFNγ/IL10 threshold, lactate, norepinephrine requirement, ACTH test analysis and calculation of odds ratios ... 3
Text S3. The Hellenic Sepsis Study Group (HSSG) cohort ... 4
Text S4. Age is not a confounder ... 5
Text S5. Integration of all cohorts ... 6
Text S6. The SISPCT trial ... 6
Text S7. The Crossover study ... 7
Text S8. Restricting the analysis to patients with IFNγ and IL10 values being within the detection limits ... 7
Text S9. Relating IFNγ/IL10 to SIRS patients with and without bacteremia ... 8
Text S10. Comparing the performance to the adenocorticotropin (ACTH) stimulation test ... 8
Text S11. Interaction analysis of treatment and biomarker ... 8
Supplementary Figures ... 9
Figure S1. The workflow ... 9
Figure S2. The calibration curve for setting the threshold ... 10
Figure S3. Interaction plots of treatment and IFNγ/IL10 ... 10
Figure S4. Distributions of serum lactate in patients with high and low ratios of IFNγ/IL10 ... 11
Figure S5. IFNγ, IL10 and IFNγ/IL10 of immune cells challenged *ex vivo* with *E.coli* fragments or LPS ... 11
Supplementary Tables ... 12
Table S1. Detailed inclusion and exclusion criteria of CORTICUS, HSSG, SISPCT and the Crossover study ... 12
Table S2. Time duration between the onset of shock and the blood drawings ... 15
Table S3. Patient characteristics of the analyzed HSSG cohort (n=162) ... 15
Table S4. List of all 137 predictors ... 16
Table S5. The predictors selected during the cross-validation runs of the CORTICUS placebo arm ... 17
Table S6. Initial cytokine and blood counts of the CORTICUS *HC* and the *placebo* arm ... 17
Table S7. Results from the machine learning models using the ACTH response data ... 17
Table S8. The lactate measurements available for the CORTICUS patients ... 18
Table S9. Results from the age stratification ... 19
Table S10. Patient characteristics of the analysed SISPCT patients and the Crossover study ... 20
Table S11. Survival rates according to high and low IFNγ/IL10, restricted to patients with IFNγ or IL10 values within the detection limits ... 22
References ... 22

### Supplementary Text

### Text S1. Blood samples and serum assays of the CORTICUS Berlin sub-cohort

Blood samples were collected on day 0 (before the corticotropin test), on day 2, on the morning of day 5 (end of full dose hydrocortisone (HC) application), on day 12 (day after HC cessation), and on days 17 and 27. The short corticotropin test was performed immediately before study drug application using blood samples taken before and 60 minutes after an intravenous bolus of 0.25 mg cosyntropin (Novartis). Blood samples were stored at 4°C for three hours to avoid time imbalances between blood collection at different sites and further processing. Serum and plasma was stored at -80°C until further analysis. Heparinized and EDTA whole blood samples were used for functional assays. At the time of the CORTICUS study, soluble mediators, interleukin-(IL)-6, 8, 10, 12 p70, interferon-γ, (IFNγ), tumor necrosis factor alpha (TNF-a), soluble TNF-receptor I (sTNF-RI), soluble FAS (all BD Biosciences OptEIA (™) Set Human), and E-selectin (R&D) were measured in serum, plasma, or culture supernatant with enzyme-linked immunosorbent assay (ELISA) according to the manufacturer's instructions. This included calculating calibration and standard curves. All measurements were done in duplicate. Calculating a variation coefficient (CV) was not part of the product description. For the cytokines, EDTA plasma was used. Hydrocortisol was measured from plasma. Surface antigens of leukocytes were measured using flow cytometry, leukocytes in EDTA, thrombocytes in citrate plasma (platelet enriched), caspase/BCL2 leukocytes with heparin. Serum lactate was measured by routine blood gas analytics. Serum lactate was measured for 51 patients at day 0. Among these, 41 patients were further observed daily for 3 days.

### Text S2. Calibration of the IFNγ/IL10 threshold, lactate, norepinephrine requirement, ACTH test analysis and calculation of odds ratios

#### Calibrating the IFNγ/IL10 threshold by the death rate

In particular the death rate of the HSSG cohort was much higher than for CORTICUS and applying our marker to HSSG needed adjustment to this discrepancy. We used the CORTICUS data to establish a calibration curve for the optimal IFNγ/IL10 threshold deciding if the ratio is high or low. In the original dataset, the optimal threshold was the 39.8 percentile of all data corresponding to the optimal ratio of 0.95 of IFNγ and IL10 serum levels. This corresponds to the original death rate of 27.7%. We now generated several new datasets weighting the non-survived patients higher (or lower) mimicking a higher (or lower) overall death rate. For every newly generated dataset, we calculated again the optimal threshold. We observed a linear dependency of the threshold according to the death rate leading to the following linear model

$$threshold_{death\,rate} = 35 + 0.215 \times death\,rate\ (in\ \%) \quad (1)$$

Using this model, we set the threshold for each validation set according to their death rates. The calibration curve applied to the investigated datasets is depicted in the Figure S2.

**Regression analysis for lactate and norepinephrine requirement**

Regression analysis for lactate: To obtain the regression functions for the time courses of serum lactate in Figure 2, patient-matched changes in lactate levels were calculated for day 1, 2 and 3, relative to the baseline (day 0) for 41 patients. These patients were grouped into four sets: IFNγ/IL10 high non-HC (n=11) (Figure 2a), IFNγ/IL10 high *HC* (n=13) (Figure 2b), IFNγ/IL10 low non-HC (n=12) (Figure 2c) and IFNγ/IL10 low HC (n=5) (Figure 2d). To find out if there is a significant decreasing trend, we performed a linear regression t-test on the time series of each patient for each panel a) to d). The linear regression test: A linear model

$$y=b_0 + b_1x \quad (2)$$

was set up for all patients (for which data was available) in each group, in which y was the lactate level and x was the time point (day). The test tests if $b_1$ is not equal to zero, following a t-statistics. The data for the ratios is given in Table S8.

Regression analysis for norepinephrine requirement: To obtain the regression functions for the time courses of norepinephrine requirement in Figure 2 (e) - (h), norepinephrine requirement at day 0, 1, 2 and 3 was investigated. These patients were grouped into two sets: patients which were treated in compliance to our rule (n= 34, panel a) and d)), and patients not in compliance with our rule (n=49, panel b) and c)). To find out if there is a significant decreasing trend, we performed a linear regression t-test (as described above) on the time series of each group.

**ACTH test analysis**

Machine learning on ACTH test data was performed using the above described cross validation scheme. Two methods were applied, (1) using the same decision tree implementation as used for the discovery set, and (2) a linear discriminant analysis (LDA) employing the implementation from the *caret* package of R (1).

**Calculating the odds ratios**

Odds ratios were calculated by the following method. The ratio of survivors to non-survivors that are treated according to our decision rule was computed. The analogous ratio was computed for all patients treated oppositely to our rule. Finally, we computed the odds ratio by dividing these two ratios. Analogously to clinical studies that compare “treatment” with “no treatment”, we compared “treated in compliance with our rule” with “treated not in compliance with our rule”. The significance was calculated using a one-sided Fisher's Exact Test.

### Text S3. The Hellenic Sepsis Study Group (HSSG) cohort

The HSSG cohort represents a prospective collection of clinical data and biosamples since 2006 of patients with documented infection and at least two signs of the systemic inflammatory response syndrome (SIRS) in 45 study sites in Greece. The study protocol was approved from the Ethics Committees of all participating hospitals. Patients were enrolled after written informed consent provided by themselves or by first-degree relatives if patients were unable to consent. Patients with infection by the human immunodeficiency virus, with less than 1,000 neutrophils/mm$^3$ and with systemic intake of more than 0.3 mg/kg of equivalent prednisone the last 15 days were excluded. Peripheral blood was drawn within the first 24 hours of the advent of signs of SIRS after puncture of one peripheral vein under aseptic conditions. Blood was centrifuged and serum was shipped to the central lab located at the 4$^{th}$ Department of Internal Medicine of ATTIKON University Hospital. All enrolled patients were reclassified into infection and sepsis in 2017 using the Sepsis-3 classification criteria (2, 3). Clinical data was recorded into one case report form (CRF) that was monitored by an independent monitor. Collected information was demographics, type of infection, severity scores, biochemistry, whole blood cell counting and blood gases, microbiology, administered antibiotics, medical therapy other than antibiotics, interventions and 28-day outcome. Among available patients and biosamples in the cohort, 342 with community-acquired pneumonia and intraabdominal infections and septic shock were randomly selected and analyzed (Table S3). Among them, for those treated with low-dose HC was 50mg intravenously four times daily for six days followed by gradually tapering off. From serum of 362 patients, secreted cytokines were measured using the LEGENDplex Human Inflammation Panel (13-plex) (BioLegend) according to manufacturer´s protocol with half of the reagents volume and sample incubation time at 4°C over night. After quality control, a total of 162 eligible shock patients (HC: n=63, No HC: n=99) were selected. If only one of the cytokines (IFNγ or IL10) was below the detection limit, the value of the detection limit was taken.

For propensity score matching, first all available HSSG baseline features (3 continuous, 11 binary and 1 categorical feature) were tested for treatment bias using Fisher's exact tests. For this testing, the features were binarized as follows. For every continuous feature, 4 binary features were calculated based on their 25 percentiles. For categorical features, entry specific binary features were calculated. This, in total, led to 25 binary features. Multiple testing correction was performed employing the method by Benjamini Hochberg (4). The multiple correction did not lead to any significant ($p<0.1$) treatment confounding features. However, two features, 'history of renal disease' and 'history of chronic heart failure' showed a significant bias before multiple correction. An ad-hoc analysis was performed in order to study if the odds ratio (OR) is affected after propensity score matching based on these two features. The propensity score matching and selection of patients was performed using the 'matchIt' package in R (5) ('genetic algorithms', n=50,000 bootstrapping iterations), leading to 126 patients and OR= 2.17 (95% CI: 1.02-4.63), p = 0.032. This confirmed that the existing treatment bias in the 'history of renal disease' and the 'history of chronic heart failure' were not the contributing factors in the performance of the predictor, IFNγ/IL10.

### Text S4. Age is not a confounder

The difference in age between the patient groups treated with *HC* and *placebo* (CORTICUS) was not confounding when we statistically stratified for age.

Statistical stratification: The average age of the HC patients was 59.4 years while for the *placebo* patients it was 69.4 years. Therefore, we tested if stratification changed our results. Stratification was performed as follows. The age of the youngest patient of the *placebo* group was 43 years. This patient was counted at 100%. For each year a *placebo* patient was older than 43 years we decreased the percentage of counting this patient by *ΔP*. In turn, the oldest *HC* patient was 87. This was fully counted in the *HC* arm and we decreased the percentage of being counted for all *HC* patients by the same amount *ΔP* for each year they are younger than 87 years. The value *ΔP* was computed in such a way that the average age in both the *HC* and the *placebo* groups equaled. The resulting confounder adjustment did not significantly alter the proposed stratification results, rounding the patient weights, yielded an odds ratio of OR=4.98 (95%CI: 1.17-21.24, p=0.02) of the survival in the responders (Table S9).

### Text S5. Integration of all cohorts

The CORTICUS (discovery cohort), HSSG, SISPCT and the Crossover study cohorts (validation cohorts) were integrated (summed up). Based on the dataset dependent threshold and the treatment, patients were grouped into four classes: low-ratio non-HC, low-ratio *HC*, high-ratio non-HC and high-ratio HC. The number of survivors and non-survivors in the groups of patients treated in compliance with our decision rule (i.e. HC treated patients with low-ratios and non-HC-treated patients with high-ratios) was compared with the number of survivors/non-survivors in the overall cohort and an odds ratio was calculated. The contingency table is shown in Table 2h. The significance was calculated using a one-sided Fisher's Exact Test.

### Text S6. The SISPCT trial

The *placebo*-controlled, randomized trial of Sodium Selenite and Procalcitonin guided antimicrobial therapy in Severe Sepsis (SISPCT) was performed in 33 intensive care units in Germany. The purpose of this study was to determine whether the intravenous application of sodium-selenite can reduce mortality in patients with severe sepsis or septic shock. Additionally, it was investigated, whether the measurement of procalcitonin - a marker of infection - can be used to guide antimicrobial therapy during the disease course. Between November 2009 and March 2013, 8,174 patients with septic shock or severe sepsis were screened and 1,089 eligible patients with informed consent were randomized. Among these, 109 patients (n=59 selenium treated and n=50 *placebo*) were included in the Munich (Ludwig-Maximilians-University, LMU) sub-study for which we performed cytokine measurements of the blood samples (ethics votum amendment EudraCT: 2007–004333-42). For our study, we excluded the selenium treated patients, and 1 patient who died at the day of inclusion. Patient characteristics are given in Table S10a. Secreted cytokine levels in blood serum samples were measured by using the LEGENDplex Human Inflammation Panel (13-plex) (BioLegend) according to manufacturer´s protocol with half of the reagents volume and sample incubation time at 4°C overnight.

For propensity score matching, first all available SISPCT baseline features (4 continuous, 20 binary and 1 categorical feature) were tested for treatment bias using Fisher's exact tests. For this testing, the features were binarized as follows. For every continuous feature, 4 binary features were calculated based on their

25 percentiles. For categorical features, entry specific binary features were calculated. This, in total, led to 40 binary features. Multiple testing correction was performed employing the method by Benjamini Hochberg (4). Significant features ($p<0.1$) were used for propensity score matching comprising ‘serum lactate’, ‘norepinephrine dosage’, ‘age’, ‘presence of septic shock (according to ACCP/SCCM criteria)’, ‘presence of severe sepsis (according to ACCP/SCCM criteria)’, ‘kidney dysfunction’ and ‘application of inotrope/pressor drug’. Finally, for propensity score matching and selection of patients, we applied the ‘matchIt’ package in R (5) (‘genetic algorithms’, n=50,000 bootstrapping iterations), leading to 24 patients with well matching propensities (Table S10b).

### Text S7. The Crossover study

The study was published elsewhere (6). Briefly, a double-blinded, randomized, *placebo*-controlled, Crossover study was performed with 40 patients diagnosed with septic shock. Until day 3, one arm received first 100 mg of HC as a loading dose and 10 mg per hour until day 3 (n=20), followed by 3 days *placebo*. The other arm received the first three days *placebo* (n=20), followed by *HC* until day 6. Blood samples were collected on day 0 (before randomization), and every subsequent day until day 6. To exclude the time point of HC application as an additional variable, we didn't regard the patients treated at day 3-5 with HC as they were neither treated as the placebo nor the verum arm in CORTICUS. Patient characteristics are given in Table S10c. In contrast, the patients treated at day 0 to day 2 were treated comparably to the CORTICUS verum arm (treated for 3 days, CORTICUS: treated for 5 days, directly at start of the study) and were used for our study. At the time of the study, serum cortisol was measured with solid-phase radioimmunoassay (Biermann, Bad Nauheim, Germany). Enzyme-linked immunosorbent assays (ELISA) were used for measurement of interleukin 4, 8, 10, 12 p70, and IFNγ (BD PharMingen, Germany), soluble E-selectin (BenderMed Alexis, Austria), IL6 (R&D, Wiesbaden, Germany), soluble tumor necrosis factor receptors I and II (Biosource, Germany). The study protocol for this study was approved by the institutional ethics committee.

### Text S8. Restricting the analysis to patients with IFNγ and IL10 values being within the detection limits

We inspected if the measured values for IFNγ and IL10 were beyond the detection limits and performed the analysis without these patients to find out if this issue may be a confounder of our analysis. In the Corticus subcohort, for 5 patients the IFNγ and for 7 patients the IL10 values were at or below the detection limit (detection limit for IFNγ: 2.35 pg/ml, for IL10: 3.90 pg/ml). Upon removing these patients from the analysis, n=73 patients remained. For HSSG n=28 were under the detection limit for IFNγ (detection limit for IFNγ: 2.94 pg/ml) and n=3 were under detection limit for IL10 (detection limit for IL10: 1.10 pg/ml). For SISPCT n=11 were under detection limit for IFNγ (detection limit for IFNγ: 5.61 pg/ml). For the Crossover study, all patients were within the detection limits for IFNγ and IL10. The results without these n=10 CORTICUS, n=31 HSSG and n=11 SISPCT patients were comparable to the results including these patients. All results without these patients are given in Table S11.

### Text S9. Relating IFNγ/IL10 to SIRS patients with and without bacteremia

Matera and coworkers investigated cytokine concentrations of 52 patients with diagnosis of systemic inflammatory response syndrome (SIRS) at hospital admission, of which n=28 were bacteremic. Two patients had septic shock. n=13 were non-survivors, n=39 survivors. SIRS was defined as two or more of the criteria 1) hypothermia or fever, 2) tachycardia; 3) tachypnea, and 4) leukocytosis, leukopenia or immature band forms (details, see (7)). Subjects under the age of 18 and patients treated with immunosuppressive drugs were excluded from the study. We obtained their cytokine measures from their article and calculated error estimates for IFNγ and IL10 concentrations with which we calculated the ratio IFNγ/IL10 and the estimated error. Error estimates for the ratio were calculated according to the Gaussian error propagation law,

$$\frac{\Delta y}{y} = \sqrt{\left(\frac{\Delta x_1}{x_1}\right)^2 + \left(\frac{\Delta x_2}{x_2}\right)^2} \qquad (3)$$

for $\Delta y$ being the error of $y(x_1, x_2)$ = IFNγ/IL10 and $x_1$=IFNγ, $x_2$= IL10.

### Text S10. Comparing the performance to the adenocorticotropin (ACTH) stimulation test

We tested two separate strategies to improve the predictive value of ACTH tests. The first strategy involved implementation of decision trees (stumps) for the ACTH stimulation response and serum cortisone baseline levels, and the other strategy involved linear discriminant (LD) models for the baseline cortisol levels in combination to the ACTH stimulation response. We followed a similar strategy as before where models were trained on *placebo* patients in a leave one out cross validation setting and the consensus model was used for getting the cross-performance in the *HC* arm. Both approaches reproduced a near-random performance quality on the data, similar as originally stated by Sprung *et al*. (8). The performance matrices with stumps and LD approaches are given in Table S12. In summary, we couldn't stratify HC therapy using the ACTH test baseline and/or response predictors.

### Text S11. Interaction analysis of treatment and biomarker

We carried out a two-way ANOVA analysis in R. A linear model for ANOVA was calculated using the *'lm'* function. 28-day survival was the independent variable. Treatment and the IFNγ/IL10 ratio were the dependent variables. An F-test was applied to assess the significance of the interaction coefficient. In the discovery set (n=83), a significant interaction (p=0.0083) between the identified biomarker and the treatment in predicting the outcome was observed. The validation set HSSG showed a similar tendency of interaction (i.e. the IFNγ/IL10 levels increase as the patients survive in the non-HC arm and *vice versa*). However, this was not significant (p = 0.13). The interaction plots are shown in Figure S3. For the data from SISPCT and the Crossover study, the test could not be applied due to the number of non-HC patients that died were zero (SISPCT), and the fact that only one arm could be analyzed (Crossover).

### Supplementary Figures

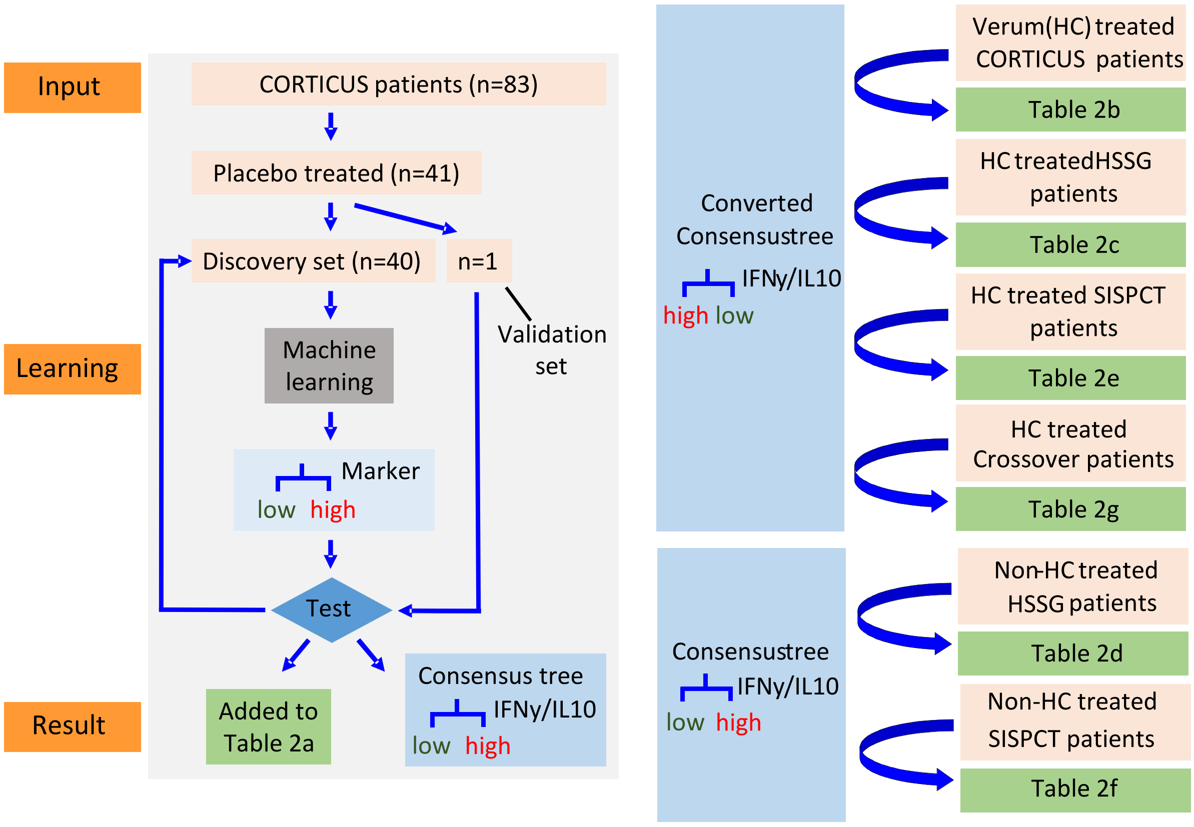


a) b)

### Figure S1. The workflow

a) The algorithm for the discovery of the theranostic marker, to note the term "validation set" is here used within the context of the cross-validation scheme:

1) From all investigated CORTICUS patients, the placebo treated patients are selected.

2) The selected patients are split into a training set (n=40 and a validation set (n=1).

3) Machine learning: Selection of the best predictor out of 137 available predictors to predict survival on the training set, using one-predictor based decision trees.

4) Testing the performance of the selected predictor on the validation set.

5) Adding the result from 4) to the confusion matrix, and storing the tree.

6) Going back to 2). In 2) the next patient is forming the validation set, and the rest of placebo patients are the training set

7) From all stored trees, a consensus tree is determined (the one which has been used most often, i.e. high IFNγ/IL10 predicts survival, low IFNγ/IL10 predicts non-survival)

b) The converted consensus tree (low IFNγ/IL10 predicts survival, high IFNγ/IL10 predicts non-survival) is applied to the HC treated patients of CORTICUS, HSSG, SISPCT, and to the early arm of the Crossover study. The consensus tree is applied to the non-HC treated patients of HSSG and SISPCT.

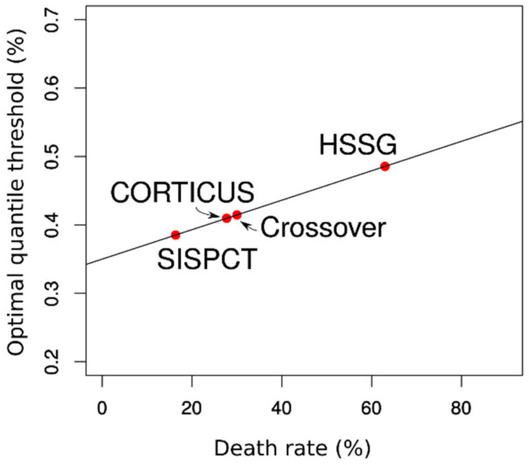


### Figure S2. The calibration curve for setting the threshold

The calibration curve is shown to set the percentile threshold of the predictor, depending on the death rate of the study. The positive gradient indicates that as the death rate increases more patients are needed to be treated with HC to obtain better survival. HSSG had a death rate of 62.96%, hence the corresponding optimal threshold was the 48.55 percentile. For SISPCT and the crossover study, the death rates were 16.33% and 30% corresponding to the 38.51 and 41.50 percentile, respectively.

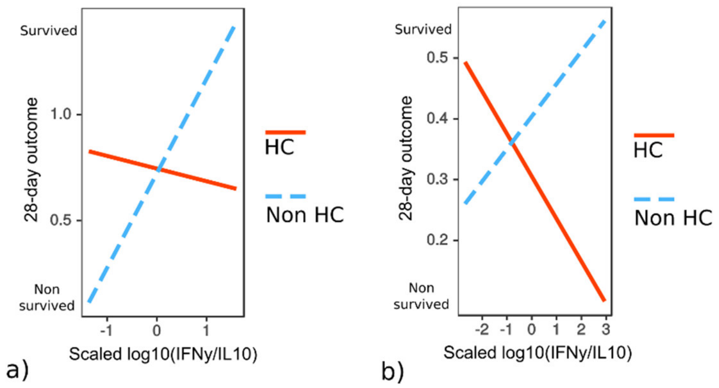


### Figure S3. Interaction plots of treatment and IFNγ/IL10

Interaction plot of treatment and IFNγ/IL10 in (a) the discovery set, CORTICUS and, (b) in the validation set, HSSG. The discovery set showed a significant interaction (p=0.0083) between treatment and the IFNγ/IL10 ratio. HSSG showed a similar tendency, but not significant (p = 0.13). Details are given in Text S9.

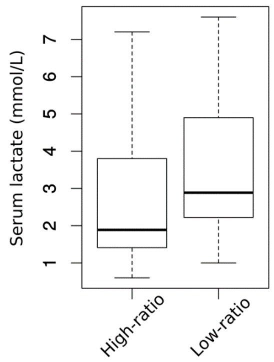


### Figure S4. Distributions of serum lactate in patients with high and low ratios of IFNγ/IL10

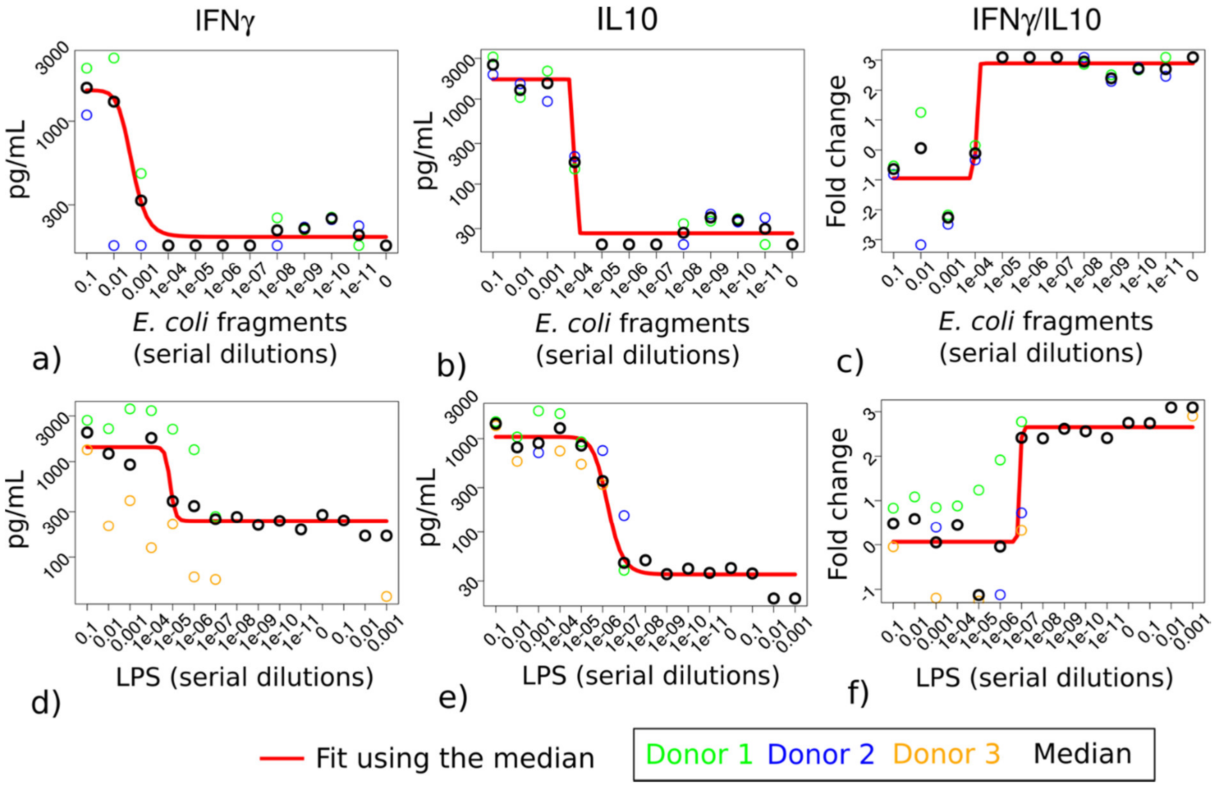


### Figure S5. IFNγ, IL10 and IFNγ/IL10 of immune cells challenged *ex vivo* with *E.coli* fragments or LPS

Whole blood from healthy donors was challenged with indicated serial dilution of a sonicated and heat inactivated *E. coli* fragments stock of a second clinical isolate (results from the first clinical isolate are shown in the main text) or LPS concentrations (0.1 ng/ml to 1mg/ml) mimicking the immunologic loads. IFNγ and IL10 levels were measured 18h after immune induction in the supernatant. The responses are consistent with the results of the first clinical isolate displayed in the main text. Both IFNγ (a, d) and IL10 (b, e) were elevated with increasing load while IFNγ/IL10 (c, f) showed the opposite behaviour, i.e. a higher load was associated with a lower ratio.

### Supplementary Tables

### Table S1. Detailed inclusion and exclusion criteria of CORTICUS, HSSG, SISPCT and the Crossover study

Taken from (3, 6, 8, 9)

| **Inclusion criteria:** | |
|---|---|
| CORTICUS | 1. Clinical evidence of infection within the previous 72 hours (may be present longer than 72 hours), only one of a, b, c, or d required:<br>a) Presence of polymorphonuclear cells in a normally sterile body fluid (excluding blood);<br>b) Positive culture or Gram staining of blood, sputum, urine or normal sterile body fluid;<br>c) Focus of infection identified by visual inspection (e.g. ruptured bowel with the presence of free air or bowel contents in the abdomen found at the time of surgery, wound with purulent drainage);<br>d) Other clinical evidence of infection - treated community acquired pneumonia, purpura fulminans, necrotising fascitis, etc. |
| | 2. Evidence of a systemic response to infection as defined by the presence of two or more of the following signs within the previous 24 hours (these signs may be present longer than 72 hours):<br>a) Fever (temperature >38.3°C) or hypothermia (rectal temperature < 35.6°C);<br>b) Tachycardia (heart rate of >90 beat/min);<br>c) Tachypnea (respiratory rate > 20 breaths/min, PaC02<32 mmHg) or patient requires invasive mechanical ventilation;<br>d) Alteration of the WBC count: >12,000 cells/$mm^3$, <4,000 cells/$mm^3$ or >10% immature neutrophils (bands). |
| | 3. Evidence of shock defined by (a and b both required within the previous 72 hours (may NOT be present longer than 72 hours).<br>a) A systolic blood pressure < 90 mmHg or a decrease in SBP of more than 50 mmHg from baseline in previous hypertensive patients (for at least one hour) despite adequate fluid replacement OR need for vasopressors for at least one hour (infusion of dopamine ≥ 5 mcg/kg/min or any dose of adrenaline, noradrenaline, phenylephrine or vasopressin) to maintain a SBP ≥ 90 mmHg;<br>b) Hypoperfusion or organ dysfunction which is not the result of underlying diseases or drugs, but is attributable to sepsis, including one of the following:<br>1. Sustained oliguria (urine output < 0.5 ml/kg/hr for a minimum of 1 hour)<br>2. Metabolic acidosis [pH of < 7.3, or a base deficit of ≥ 5.0 mmol/L, or an increased lactic acid concentration (> 2 mmol/L)].<br>3. Arterial hypoxemia (Pa02/FI02<280 in the absence of pneumonia)(Pa02/FI02<200 in the presence of pneumonia).<br>4. Thrombocytopenia - platelet count ≤ 100,000 cells/$mm^3$.<br>5. Acute altered mental status (Glasgow Coma Scale < 14 or acute change from baseline). |
| | 4. Age >= 18 years |
| | 5. Informed Consent |
| | 6. Measured cortisol level at baseline and 60 minutes after 0.25 mg cosyntropin stimulation |
| HSSG | 1. Patients reclassified into infection and sepsis using the Sepsis-3 classification criteria (2, 3) |
| | 2. Age >= 18 years |
| | 3. Informed Consent |
| | 4. Community-acquired pneumonia and intraabdominal infections |
| SISPCT | 1. Onset of severe sepsis (A and B both required) or septic shock (C) <24 h. Severe sepsis was defined as the presence of microbiologically proven, clinically proven, or suspected |

| | |
|---|---|
| | infection; presence of Systemic Inflammatory Response Syndrome (A); and development of at least one organ dysfunction (B) within the last 24 hours.<br><br>A) Diagnosis of SIRS required the fulfilment of at least two of the following criteria: hypo- (≤36°C) or hyperthermia (≥38°C), tachycardia (≥90 bpm); tachypnea (≥20 breaths/min) and/or an arterial pCO2 ≤4.3 kPa (32 mmHg) and/or mechanical ventilation; leukocytosis ≥12000/µl or leukopenia ≤4000/µl and/or a left shift in the differential white blood cell count ≥10%<br>B) For the diagnosis of organ dysfunction one of the following criteria had to be fulfilled:<br>i. Presence of acute encephalopathy with reduced vigilance, agitation, disorientation, delirium not explained by psychotropic medication,<br>ii. Thrombocytopenia ≤100.000/µl or a drop in the thrombocyte count >30% within 24 hours not explained by hemorrhage,<br>iii. Arterial hypoxemia with an arterial pO2 <10 kPa (75 mmHg) when breathing normally or an oxygenation index (paO2/FiO2 ≤33kPa (250 mmHg) not explained by presence of a pulmonary or cardiac disease,<br>iv. Arterial hypotension with a systolic blood pressure ≤90 mmHg or mean arterial blood pressure ≤70 mmHg for at least one hour despite adequate fluid loading not explained by other causes of shock,<br>v. Renal dysfunction with an urine output ≤0.5 ml/kg/h for at least one hour despite adequate fluid loading and/or increase of serum creatinine more than twofold above the reference range of the local laboratory,<br>vi. Metabolic acidosis with a base deficit ≥5.0 mmol/l or a serum lactate ≥1.5 fold above the reference range of the local laboratory.<br>C) Septic shock was defined as the presence of infection and SIRS as defined for severe sepsis as well as presence of arterial hypotension with a systolic blood pressure ≤90 mmHg or a mean arterial blood pressure ≤70 mmHg for at least 2 hours or administration of a vasopressor (dopamin ≥5 µg kg-1 min-1; norepinephrine, epinephrine, phenylephrine, or vasopressin in any dosage) to maintain systolic blood pressure ≥90 mmHg or mean arterial blood pressure ≥70 mmHg despite adequate fluid loading. |
| | 4. Age >= 18 years |
| | 5. Informed consent |
| Crossover | 1. Presence of septic shock including,<br>a) Proven or strongly suspected infection<br>b) Three or more of these conditions: mechanical ventilation, heart rate of more than 90 beats per minute, temperature of more than 38°C or less than 36°C, a white blood cell count of more than 12,000 cells/µl or less than 4,000 cells/µl, or more than 10% immature cells<br>c) Sepsis-induced hypotension (systolic blood pressure of less than 90 mm Hg or a reduction of more than 40 mm Hg from baseline in the absence of other causes of hypotension) |
| | 2. Patients requiring norepinephrine to maintain a mean arterial pressure of more than 70 mm Hg despite adequate fluid resuscitation. |
| | 2. Age >= 18 years |
| | 3. Informed Consent |
| **Exclusion criteria:** | |
| CORTICUS | 1. Pregnancy,<br>2. Age less than 18,<br>3. Underlying disease with a prognosis for survival of less than 3 months, |

| | |
|---|---|
| | 4. Cardiopulmonary resuscitation within 72 hours before study,<br>5. Drug-induced immunosuppression, including chemotherapy or radiation therapy within 4 weeks before the study,<br>6. Administration of chronic corticosteroids in the last 6 months or acute steroid therapy (any dose) within 4 weeks (including inhaled steroids). Topical steroids are not exclusions,<br>7. HIV positivity,<br>8. Presence of an advanced directive to withhold or withdraw life sustaining treatment (i.e. DNR),<br>9. Advanced cancer with a life expectancy less than 3 months,<br>10. Acute myocardial infarction or pulmonary embolus,<br>11. Another experimental drug study within the last 30 days,<br>12. Moribund patients likely to die within 24 hours,<br>13. Patients in the ICU for more than 2 months at the time of the start of septic shock, |
| HSSG | 1. Infection by the human immunodeficiency virus,<br>2. < 1,000 neutrophils/$mm^3$<br>3. Systemic intake of more than 0.3mg/kg of equivalent prednisolone the last 15 days |
| SISPCT | 1. Pregnant or breast-feeding women,<br>2. Fertile female women without effective contraception,<br>3. Participation in interventional clinical trial within the last 30 days,<br>4. Current participation in any study,<br>5. Former participation in this trial,<br>6. Selenium intoxication,<br>7. No commitment to full patient support (i.e. DNR order),<br>8. Patient's death is considered imminent due to coexisting disease,<br>9. Relationship of the patient to study team member (i.e. colleague, relative),<br>10. Infection where guidelines recommend a longer duration of antimicrobial therapy (i.e. endocarditis, tuberculosis, malaria etc),<br>11. Immunocompromised patients. |
| Crossover | 1. Pregnancy,<br>2. Glucocorticoid medication within the last 3 months,<br>3. Ongoing immunosuppressive therapy,<br>4. Hematologic diseases,<br>5. Moribund state. |

### Table S2. Time duration between the onset of shock and the blood drawings

| Pat. Code | Time to HC (h) | Pat. Code | Time to HC (h) | Pat. Code | Time to HC (h) | Pat. Code | Time to HC (h) |
|---|---|---|---|---|---|---|---|
| 660261 | 20 | 610242 | 59 | 940373 | 24 | 820328 | 53.5 |
| 670265 | 3 | 600240 | 61 | 850337 | 26 | 920368 | 35 |
| 660262 | 16 | 680270 | 14.5 | 900357 | 21 | 1370546 | 43 |
| 660264 | 28.7 | 690276 | 19 | 900359 | 30 | 1370545 | 37.5 |
| 660263 | 18 | 850339 | 49 | 890353 | 23 | 840335 | 66.5 |
| 670266 | 19 | 610241 | 47 | 570225 | 20 | 840336 | 71 |
| 620245 | 7 | 850340 | 19 | 810323 | 30 | 870345 | 7 |
| 570226 | 41.5 | 630250 | 6 | 890354 | 8.5 | 870346 | 17.5 |
| 650258 | 29 | 920367 | 44 | 800319 | 29 | 820325 | 67 |
| 620246 | 26 | 880350 | 24 | 900360 | 5 | 910363 | 27.5 |
| 650257 | 19 | 920366 | 32 | 600239 | 25 | 1370547 | 23 |
| 620247 | 29 | 610244 | 15 | 810324 | 12 | **Mean±sd** | **29.4±16.6** |
| 620248 | 58 | 650259 | 16 | 800320 | 23 | | |
| 650260 | 41 | 590235 | 24 | 940374 | 18 | | |
| 580229 | 13.5 | 680269 | 39 | 840334 | 39 | | |
| 580232 | 24.5 | 880352 | 23 | 840333 | 21 | | |
| 690273 | 5 | 810321 | 21 | 930372 | 24 | | |
| 590234 | 1.45 | 590236 | 28 | 880349 | 30 | | |
| 850338 | 45 | 630249 | 24.5 | 880351 | 30 | | |
| 580231 | 49 | 800317 | 13 | 910361 | 41 | | |
| 920365 | 56 | 900358 | 42 | 820327 | 53.5 | | |
| 690275 | 18 | 610243 | 25.5 | 570227 | 51 | | |
| 670268 | 67.5 | 800318 | 26.5 | 570228 | 51 | | |
| 590233 | 7 | 810322 | 23 | 860342 | 44 | | |

### Table S3. Patient characteristics of the analyzed HSSG cohort (n=162)

| HSSG | Non HC (n=99) | HC (n=63) |
|---|---|---|
| Gender (male/ female, n) | 38/61 | 32/31 |
| Age, years (mean, 95% CI) | 73.10 (70.29-75.81) | 71.10 (67.41-74.78) |
| SOFA | 9.03 (8.33-9.73) | 8.83 (7.95-9.70) |
| APACHE II | 24.46 (22.89-26.04) | 22.67 (20.91-24.42) |
| 28 days survival (n, %) | 40 (40%) | 20 (32%) |
| Site of infection (n, %) | | |
| - Community-acquired pneumonia | 62 (63%) | 33 (52%) |
| - Intrabdominal infection | 37 (37%) | 30 (48%) |
| Co-morbidities (n, %) | | |
| - Presence of acute kidney injury | 25 (25%) | 22 (35%) |
| - History of diabetes mellitus type 2 | 26 (26%) | 21 (33%) |
| - History of renal disease | 16 (16%) | 3 (5%) |
| - History of chronic heart failure | 34 (34%) | 11 (17%) |
| - History of chronic obstructive pulmonary disease | 19 (19%) | 11 (17%) |

### Table S4. List of all 137 predictors

| | | | |
|---|---|---|---|
| 42/14 Mean | IL10 (Con-A) [pg/ml] | Natural killer cells [%] | SOFA Resp |
| 42/14 UR% | IL10 (LPS) [pg per 1000 cells] | Natural killer cells [1/nl] | T lymphocytes [%] |
| 42/14ADP M | IL10 (LPS) [pg/ml] | Nitrit/Nitrat [µmol/l] | T lymphocytes [1/nl] |
| 42/14ADP UR | IL10 [pg/ml] | Norepinephrine | T-Helper lymphocytes[%] |
| Age | IL10/TNFα ratio | OSF*_card | T-Helper lymphocytes[1/nl] |
| B lymphocytes [%] | IL10/TNFα ratio (LPS) | OSF_coag | T-Suppressor lymphocytes [%] |
| B lymphocytes [1/nl] | IL12/IFNγ ratio | OSF_liv | T-Suppressor lymphocytes [1/nl] |
| BE high | IL12/IL10 ratio | OSF_nerv | Temperature |
| BE low | IL12/TNFα ratio | OSF_renal | TH-Akt. Caspase 3 pos [% ] |
| Bicarbonate high | IL6 (LPS) [pg per 1000 cells] | OSF_resp | TH-Akt. Caspase 3 pos [1/nl] |
| Bicarbonate low | IL6 (LPS) [pg/ml] | PaCO2 high | TH-BCL-2 pos [% gated] |
| Bilirubin total high | IL6 [pg/ml] | PaCO2 low | TH-BCL-2 pos [1/nl] |
| CD11b expression on PMN (Mean) | IL6/IFNγ ratio | PaO2 low | Thrombocytes [1/nl] |
| Creatinine high | IL6/IFNγ ratio (LPS) | PEEP high | Tidal volume |
| D-Dimer [µg/ml] | IL6/IL10 ratio | pH high | Tidal volume [/kg] |
| DIC-Overall score | IL6/IL10 ratio (LPS) | pH low | TNFα (LPS) [pg per 1000 cells] |
| Dobutamine | IL6/IL12 ratio | Platelets high | TNFα (LPS) [pg/ml] |
| E-Selectin [pg/ml} | IL6/IL8 ratio | Platelets low | TNFR1 |
| Factor VII [%] | IL6/TNFα ratio | PMNs [%] | TPZ [sec.] |
| FiO2 high | IL6/TNFα ratio (LPS) | PMNs [1/nl] | TS-Akt. Caspase 3 pos [%] |
| GCS | IL8 [pg/ml] | Protein C [%] | TS-Akt. Caspase 3 pos [1/nl] |
| Gender | IL8/IFNγ ratio | PTT high | TS-BCL2 pos [%] |
| HbO2 low | IL8/IL10 ratio | Respiratory rate | TS-BCL2 pos [1/nl] |
| Heart rate | IL8/IL12 ratio | SBP | Urea high |
| Hemoglobin high | IL8/TNFα ratio | Score D-Dim | Urinary output |
| HLA-DR expression on monocytes (Mean) | IL12 [pg/ml] | Score Thr | WBC high |
| HLA-DR-receptors on monocytes [1/cell] | INR | Score TPZ | WBC low |
| IFNγ (Con-A) [pg per 1000 cells] | Lactate high | Sedation | Weight |
| IFNγ (Con-A) [pg/ml] | Leukocytes [1/nl] | sFas | |
| IFNγ [pg/ml] | Lymphocytes [%] | SOFA | |
| IFNγ/IL10 ratio | Lymphocytes [1/nl] | SOFA Cardio | |

*OSF: Organ system failure

### Table S5. The predictors selected during the cross-validation runs of the CORTICUS placebo arm

| Feature | Number of times selected |
|---|---|
| IFNγ/IL10 < 0.95 | 39 |
| TNF-a (pg/ml) < 22115.78 | 2 |

### Table S6. Initial cytokine and blood counts of the CORTICUS *HC* and the *placebo* arm

| | Hydrocortisone (HC) | Placebo (PL) | p* |
|---|---|---|---|
| **A. Cytokines** | Median (Q1-Q3) | Median (Q1-Q3) | |
| IFNγ [pg/ml] | 35.05 (15.79-152.13) | 30.16 (17.93-57.80) | 0.14 |
| IL10 [pg/ml] | 31.21 (13.30-76.22) | 31.66 (15.68-54.76) | 0.63 |
| IL12 [pg/ml] | 10.03 (3.90-43.60) | 13.00 (3.90-43.27) | 0.66 |
| IL6 [pg/ml] | 388.02 (177.80-487.27) | 377.21 (237.17-509.01) | 0.61 |
| IL8 [pg/ml] | 130.95 (56.63-226.28) | 86.81 (53.81-244.44) | 0.4 |
| sFas [pg/ml] | 2137.17 (1544.84-3335.26) | 2155.38 (1551.38-3196.85) | 0.9 |
| sTNF-R1 [pg/ml] | 25806.28 (13322.69-40599.84) | 18809.52 (12009.18-28885.86) | 0.16 |
| **B. Blood counts** | | | |
| Leukocytes [/nl] | 13.35 (10.10-17.23) | 12.45 (9.13-16.93) | 0.94 |
| Lymphocytes [/nl] | 0.71 (0.34-0.93) | 0.75 (0.41-1.05) | 0.54 |
| Monocytes [/nl] | 0.53 (0.36-0.80) | 0.56 (0.33-0.92) | 0.82 |
| NK cells [/nl] | 0.04 (0.02-0.07) | 0.05 (0.03-0.12) | 0.21 |
| PMNs [/nl] | 11.69 (7.17-14.36) | 11.06 (7.75-15.21) | 0.88 |
| B-lymphocytes [/nl] | 0.09 (0.04-0.20) | 0.08 (0.05-0.13) | 0.43 |
| T-Helper lymphocytes [/nl] | 0.30 (0.13-0.43) | 0.26 (0.14-0.50) | 0.72 |
| T-Suppressor lymphocytes [/nl] | 0.10 (0.05-0.14) | 0.07 (0.04-0.17) | 0.89 |
| Thrombocytes [/nl] | 170.00 (99.50-250.00) | 146.50 (92.00-203.25) | 0.43 |

* Two-sided Wilcoxon test

### Table S7. Results from the machine learning models using the ACTH response data

| Stumps: Response to ACTH | Suggested | All patients |
|---|---|---|
| Non-survivors (predicted) | 15 | 23 |
| Survivors (predicted) | 36 | 60 |
| Survival rate | 71% | 72% |

| LD: Baseline + Response to ACTH | Suggested | All patients |
|---|---|---|
| Non-survivors (predicted) | 12 | 23 |
| Survivors (predicted) | 30 | 60 |
| Survival rate | 71% | 72% |

### Table S8. The lactate measurements available for the CORTICUS patients

| Day 0 | Day 1 | Day 2 | Day 3 | Class |
|---|---|---|---|---|
| 1.9 | 1.3 | 1.6 | 1 | PL High-ratio |
| 2.4 | 2.1 | 1.3 | 1.4 | PL High-ratio |
| 3.4 | 17 | 10 | 6 | HC High-ratio |
| 2.9 | NA | NA | NA | PL Low-ratio |
| 0.8 | 11 | 1.3 | 14 | HC High-ratio |
| 1.9 | 1.3 | 1 | 1.1 | HC High-ratio |
| 3.4 | 3.5 | 2.4 | 2.4 | HC High-ratio |
| 4.9 | 2.5 | 2.2 | 1.4 | PL Low-ratio |
| 7.2 | 2.1 | 1.4 | 1.8 | PL High-ratio |
| 4.1 | 3 | 1.2 | 1.4 | HC High-ratio |
| 0.9 | 0.3 | 0.2 | 0.1 | PL High-ratio |
| 1.4 | NA | NA | NA | HC High-ratio |
| 5.1 | 5 | 10.4 | 5.9 | PL Low-ratio |
| 2.3 | 2.5 | 1.6 | 1.4 | HC Low-ratio |
| 3.4 | 2.4 | 1.8 | 2.1 | PL Low-ratio |
| 5.5 | 1.4 | 1.6 | 1.3 | PL Low-ratio |
| 3.6 | 1.6 | 1.1 | 1.9 | HC High-ratio |
| 2.5 | NA | NA | NA | PL Low-ratio |
| 0.6 | NA | NA | NA | PL High-ratio |
| 4 | 4 | 1.2 | 1.4 | PL High-ratio |
| 7.6 | NA | NA | NA | HC Low-ratio |
| 1.7 | 2.4 | 2.2 | 1.6 | HC High-ratio |
| 3 | NA | NA | NA | PL High-ratio |
| 1.2 | NA | NA | NA | HC High-ratio |
| 4.1 | 3.8 | 2.4 | 3.1 | HC High-ratio |
| 5.2 | 2.7 | 1.6 | 1.4 | HC Low-ratio |
| 1.8 | 2.5 | 3.1 | 5.4 | HC High-ratio |
| 1.6 | 1.5 | 1.4 | 1.6 | PL High-ratio |
| 6.3 | 5 | 2.2 | 0.5 | HC High-ratio |
| 1.1 | 1.3 | 1.4 | 1 | PL High-ratio |
| 1.5 | NA | NA | NA | HC High-ratio |
| 1.7 | NA | NA | NA | PL Low-ratio |
| 4.4 | 1.2 | 1.2 | 0.8 | HC High-ratio |
| 2.2 | 2.3 | 2.6 | 1.8 | HC High-ratio |
| 1 | 0.8 | 1.1 | 1.1 | PL High-ratio |

| | | | | |
|---|---|---|---|---|
| 2.9 | 2.4 | 2.3 | 2.2 | PL Low-ratio |
| 2.7 | 2.4 | 2.2 | 2.1 | HC Low-ratio |
| 1.4 | NA | NA | NA | PL High-ratio |
| 1.7 | NA | NA | NA | HC High-ratio |
| 2 | 1.6 | 1.7 | 1 | PL Low-ratio |
| 1.4 | 1.9 | 2.2 | 3 | PL Low-ratio |
| 5 | 5 | 4.1 | 4.4 | PL Low-ratio |
| 3.6 | 2.4 | 1.5 | 1.5 | PL Low-ratio |
| 1 | 0.9 | 1 | 1 | HC Low-ratio |
| 1.6 | 1.2 | 1.1 | 0.9 | PL High-ratio |
| 1.1 | 17 | 9 | 12 | PL Low-ratio |
| 1.3 | 1.9 | 2.2 | 1.8 | HC High-ratio |
| 3.9 | NA | NA | NA | PL Low-ratio |
| 5.3 | 2.6 | 1.8 | 1.7 | PL High-ratio |
| 5.1 | 1.6 | 1.8 | 1.4 | PL High-ratio |
| 2.8 | 2.9 | 2.3 | 2 | PL Low-ratio |
| 3.7 | 2.7 | 2.1 | 2.9 | PL Low-ratio |
| 2.2 | 2.4 | 1.8 | 1.2 | HC Low-ratio |

### Table S9. Results from the age stratification

| *Placebo* | Non-survival | Survival | **% Survival** |
|---|---|---|---|
| IFNγ/IL10 high | 0.64* | 11.16 | **95%** |
| IFNγ/IL10 low | 5.40 | 6.07 | **53%** |

| HC | Non-survival | Survival | **% Survival** |
|---|---|---|---|
| IFNγ/IL10 high | 5.53 | 7.99 | **59%** |
| IFNγ/IL10 low | 1.57 | 7.94 | **83%** |

* The numbers show the age weighted outcome. The reasoning to perform this analysis was to understand if the age difference in the placebo and HC arm of CORTICUS affect the proposed theranostic based stratification. The method used to calculate these numbers is described in Text S4.

### Table S10. Patient characteristics of the analysed SISPCT patients and the Crossover study

#### a) All patients from the placebo arm (n=49)

| SISPCT | HC (n=35) | No HC (n=14) |
|---|---|---|
| Sex (male/ female, n) | 24/ 11 | 6/ 8 |
| Age, years (mean, 95% CI) | 63.5 (58.8-68.3) | 54.2 (44.4-63.9) |
| Septic shock at day 0 (n, %)* | 32 (91%) | 9 (64%) |
| Severe sepsis at day 0 (n, %)* | 0 (0%) | 4 (29%) |
| On inotropes/pressors at day 0 (n, %) | 32 (91%) | 10 (71%) |
| Norepinephrine requirement, mcg/kg/min at day 0 (mean, 95%CI) | 2.41 (1.77-3.05) | 1.04 (0.67-1.44) |
| Lactate, mmol/L at day 0 (mean, 95% CI) | 4.89 (3.68-6.10) | 1.96 (1.60-2.32) |
| Weight, kg (mean, 95% CI) | 81.9 (74.3-89.4) | 81.62 (71.6-91.6) |
| Survival (n) | | |
| - Day 28 (survivors/ non-survivors) | 27/ 8 | 14/ 0 |
| - Day 90 (survivors/ non-survivors) | 24/ 11 | 14/ 0 |
| Underlying infection (n, %) | | |
| - Pneumonia | 19 (54%) | 9 (64%) |
| - Urogenital | 6 (17%) | 1 (7%) |
| - Abdominal | 6 (17%) | 2 (14%) |
| - Soft-tissue | 4 (12%) | 1 (7%) |
| - Unclear | 0 (0%) | 1 (7%) |

*According to the ACCP/SCCM criteria

#### b) Propensity score matched patients (n=24)

| SISPCT | HC (n=10) | No HC (n=14) |
|---|---|---|
| Sex (male/ female, n) | 7/ 3 | 6/ 8 |
| Age, years (mean, 95% CI) | 53.8 (60.6-67.5) | 54.2 (44.4-63.9) |
| Septic shock at day 0 (n, %)* | 7 (70%) | 9 (64%) |
| Severe sepsis at day 0 (n, %)* | 0 (0%) | 4 (29%) |
| On inotropes/pressors at day 0 (n, %) | 8 (80%) | 10 (71%) |
| Norepinephrine requirement, mcg/kg/min at day 0 (mean, 95%CI) | 1.28 (0.90-1.66) | 1.04 (0.67-1.44) |
| Lactate, mmol/L at day 0 (mean, 95% CI) | 2.36 (1.63-3.08) | 1.96 (1.60-2.32) |
| Weight, kg (mean, 95% CI) | 77.2 (61.1-93.4) | 81.62 (71.6-91.6) |
| Survival (n) | | |
| - Day 28 (survivors/ non-survivors) | 7/ 3 | 14/ 0 |
| - Day 90 (survivors/ non-survivors) | 7/ 3 | 14/ 0 |
| Underlying infection (n, %) | | |
| - Pneumonia | 9 (90%) | 9 (64%) |

| | | |
|---|---|---|
| - Urogenital | 0 (0%) | 1 (7%) |
| - Abdominal | 0 (0%) | 2 (14%) |
| - Soft-tissue | 1 (10%) | 1 (7%) |
| - Unclear | 0 (0%) | 1 (7%) |

* According to the ACCP/SCCM criteria

**c) Patient characteristics of the Crossover study**

| Crossover study | HC (n=20) |
|---|---|
| Age, years (mean, 95% CI) | 54 (46, 63) |
| Sex (male/female, n) | 13/7 |
| SAPS II | 42 (35, 49) |
| SOFA | 9.7 (8.5, 10.9) |
| 28 day survival (n, %) | 14 (70%) |
| Time between onset of septic shock and inclusion, hours (n, %) | |
| - < 24 | 4 (20%) |
| - 24–48 | 7 (35%) |
| - 48–120 | 6 (30%) |
| - > 120 | 3 (15%) |
| Main source of infection (n, %) | |
| - Pulmonary | 12 (60%) |
| - Gastrointestinal | 8 (40%) |
| Microbiology | |
| - Gram positive | 3 (15%) |
| - Gram negative | 10 (50%) |
| - Mixed | 3 (15%) |
| - Fungal | 1 (5%) |
| - Not identified | 3 (15%) |

### Table S11. Survival rates according to high and low IFNγ/IL10, restricted to patients with IFNγ or IL10 values within the detection limits

a) CORTICUS patients treated with *placebo*

| | Non-survivors | Survivors | % Survivors |
|---|---|---|---|
| IFNγ/IL10 high | 3 | 18 | **86 %** |
| IFNγ/IL10 low | 7 | 8 | **53 %** |

b) CORTICUS patients treated with HC

| | Non-survivors | Survivors | % Survivors |
|---|---|---|---|
| IFNγ/IL10 high | 8 | 18 | **69 %** |
| IFNγ/IL10 low | 2 | 9 | **82 %** |

c) HSSG patients not treated with HC

| | Non-survivors | Survivors | % Survivors |
|---|---|---|---|
| IFNγ/IL10 high | 22 | 21 | **49 %** |
| IFNγ/IL10 low | 27 | 10 | **27 %** |

d) HSSG patients treated with HC

| | Non-survivors | Survivors | % Survivors |
|---|---|---|---|
| IFNγ/IL10 high | 18 | 6 | **25 %** |
| IFNγ/IL10 low | 19 | 8 | **30 %** |
